# Food quality and *vgll3* genotype influence reproductive traits in female Atlantic salmon

**DOI:** 10.1101/2024.12.12.628099

**Authors:** Katja Susanna Maamela, Eirik Ryvoll Åsheim, Ronan James O’Sullivan, Paul Vincent Debes, Andrew Herbert House, Petra Liljeström, Jenni Maria Prokkola, Petri Toivo Niemelä, Jaakko Erkinaro, Kenyon Brice Mobley, Craig Robert Primmer

## Abstract

Age at maturity is an important factor contributing to life-history diversity. In Atlantic salmon, this trait often shows sex-specific variation, but female salmon are seldom included in experimental studies of maturation. As a result, there is a gap in our knowledge of how different genetic and environmental factors affect female maturation. Using a 4-year common-garden experiment, we assessed the influence of diet (low-fat vs. control) and *vgll3* (a candidate gene in a genomic region known to influence age at maturity in Atlantic salmon) on maturation and related phenotypic traits of female Atlantic salmon from two 2^nd-^generation hatchery populations. We found the early-maturation-associated *vgll3\**E allele to be associated with higher probability of maturation. Heritability of maturation was estimated to be 0.295 with *vgll3*’s contribution to phenotypic variance being ∼2%. In addition, both body size and body condition measured in the spring prior to spawning influenced maturation. Body condition, in turn, was influenced by population and diet. The northern Oulu population and the low-fat diet were associated with lower body condition compared to the southern Neva population and the control diet. Moreover, there was an interaction between population and diet on body condition, suggesting that populations may respond differently to nutrient availability. These results broaden our understanding of the processes underlying maturation and demonstrate that genes and environment interact to shape age at maturity in female Atlantic salmon.

## Introduction

Understanding the processes governing life-history trait variation is at the forefront of evolutionary biology research (Stearns, 2000). The rapid development of genomic tools over the past two decades has allowed us to gain a fine-scale understanding of the genetic basis of life-history traits in various taxa at single-gene and genomic-region levels. For example, specific genomic regions are known to explain variation in reproductive strategies in ruffs (Küpper et al., 2016; Lamichhaney et al., 2016), developmental time in *Tribolium* beetles (Cheng et al., 2024), colouration and life-history strategy in *Colias* butterflies (Woronik et al., 2019), and migration timing and age at maturity in some salmonid species (Ayllon et al., 2015; Barson et al., 2015; Hess et al., 2016; Prince et al., 2017). These studies have expanded our knowledge of the genetic basis of life-history traits and offer an opportunity for follow-up studies that look beyond the genetic associations and into how the genomic regions or candidate genes mediate traits independently or via interactions with environmental factors.

### Atlantic salmon life-history, age at maturity, and *vgll3*

Atlantic salmon, *Salmo salar*, is one of the species for which we have gained a deeper understanding of the genetic basis of life-history trait variation in recent years, especially so with respect to age at maturity (Ayllon et al., 2015; Barson et al., 2015; Sinclair-Waters et al., 2020). Atlantic salmon have a complex life cycle often spanning both freshwater and marine environments (Jonsson & Jonsson, 2011; Thorstad et al., 2010). Consequently, they exhibit great variation in different life-history traits, such as time spent in freshwater or at sea, age or size at maturity, and number of reproductive events (de Eyto et al., 2015; Erkinaro et al., 2019; Mobley et al., 2020; Persson et al., 2023). Age at maturity is a key life-history trait tightly linked with fitness that is affected by both environment and genetics (Mobley et al., 2021) in which it has both a strong single-gene (Ayllon et al., 2015; Barson et al., 2015) and polygenic component (Sinclair-Waters et al., 2020). Allelic variation in the primary candidate gene *vgll3*, explains close to 40 percent of variation in sea age at maturity in Northern European Atlantic salmon populations (Barson et al., 2015). In relation to age at maturity, *vgll3* has two alleles, the E and the L allele, associated with early and late maturation, respectively (Barson et al., 2015). In salmonids, age at maturity often shows sex-specific variation where, on average, females mature at an older age and larger body size compared to males (e.g., Erkinaro et al., 2019; Evans et al., 2019; Fleming & Einum, 2011; Mobley et al., 2020, 2024; Tréhin et al., 2021). Further, patterns of sex-specific dominance for age at maturity at the *vgll3* locus have been reported in some wild populations with the heterozygous EL males maturing at a younger age compared to heterozygous females (Barson et al., 2015; Besnier et al., 2024; Czorlich et al., 2018; Miettinen et al., 2024; Mobley et al., 2024; Raunsgard et al., 2023). For female salmon, body size is an important determinant of reproductive fitness as larger body size allows for higher fecundity and may be associated with a better ability to dig and defend nests at the spawning grounds (Fleming, 1996). Thus, older age at maturity offers a fitness benefit, but simultaneously there is a trade-off as a longer marine migration is associated with a higher risk of mortality (Mobley et al., 2021).

Various environmental factors have a strong influence on age at maturity in salmon. For example, temperature and diet can influence growth and available energy reserves and thereby, age at maturity (Jonsson & Jonsson, 2011; Taranger et al., 2010). Maturation is an energy-demanding process (Ginther et al., 2024) and larger energy reserves may be associated with a higher probability of maturation (Bernardo, 1993; Rowe et al., 1991; Shearer & Swanson, 2000; Taranger et al., 2010). Salmon allocate almost 60 percent of their energy stores to reproduction. In female salmon, a major part of that energy is allocated to egg production (Fleming & Einum, 2011). Therefore, the ability of individual salmon to acquire energy, be it through the quantity or energetic quality of their feed, can influence growth and consequently size and age at maturity (Jonsson et al., 2012, 2013; Vollset et al., 2022).

Since the discovery of the association between *vgll3* and age at maturity, a number of studies have investigated how *vgll3* relates to different life-history traits in Atlantic salmon with a view to understanding the mechanisms behind the gene’s association with maturation. The association between *vgll3* and sea age at maturity has been demonstrated in wild populations for both males and females (Czorlich et al., 2018; Miettinen et al., 2024; Mobley et al., 2024; Raunsgard et al., 2023), and replicated for males in common-garden experiments in hatchery populations (Åsheim et al., 2023; Debes et al., 2021) and in an aquaculture strain (Ayllon et al., 2019; Sinclair-Waters et al., 2022). In addition to age at maturity, *vgll3* has been associated with behavioural traits (Bangura et al., 2022, 2024), body condition (Debes et al., 2021; House et al., 2023), migration activity (Niemelä et al., 2022), and aerobic scope (Prokkola et al., 2022). Though the link between *vgll3* and age at maturity has been studied in females in wild populations (Barson et al., 2015; Czorlich et al., 2018; Miettinen et al., 2024; Mobley et al., 2024; Raunsgard et al., 2023), only one previous study (Ayllon et al., 2019) has investigated *vgll3*’s role in female maturation in a controlled common-garden experiments where the rearing environment of the fish could be modified. In that study, no association between *vgll3* genotype and maturation age in the Mowi aquaculture strain was found (Ayllon et al., 2019). Therefore, there is an incomplete understanding of how *vgll3* and environment influence age at maturity in female Atlantic salmon.

### Objectives

In this study, we conducted a common-garden experiment to investigate how environment, namely energy content of the diet, and variation in *vgll3* influence maturation and reproductive traits in four-year-old female Atlantic salmon. With this experiment, we aimed to answer the following questions: 1) Does *vgll3* influence age at maturity in female Atlantic salmon and is the potential influence similar to that reported in wild populations? 2) does diet energy content influence maturation? 3) are there population-specific differences in maturation response to diet or in how *vgll3* genotype influences maturation? In addition to maturation, we investigate the effect of diet, population, and *vgll3* on maturation-related traits, such as body condition, body size, and fecundity to assess how these traits are affected by energy resources and genetics. We also estimate heritability of maturation and the contribution of *vgll3* to phenotypic variance to better understand the potential for selection to shape age at maturity.

## 2.0 Methods

### 2.1. Populations and cross-design

Detailed descriptions of the Atlantic salmon hatchery populations used in this study, the crossing design, and rearing conditions are provided in Åsheim et al., (2023) and description of the egg incubation phase in Debes et al., (2020), 2021). Briefly, the Atlantic salmon used in this study represent two genetically distinct Atlantic salmon populations from the Baltic Sea: the Neva and the Oulu populations, named after their locations of origin (Säisä et al., 2005). Broodstocks originating from these populations are maintained by the Natural Resources Institute Finland (Luke) for compensatory stocking purposes in Finnish rivers with hydropower regulation. The southern Neva population is routinely stocked into river Kymijoki (60.48°N, 26.89°E), which drains into the Gulf of Finland, whereas the northern Oulu population is stocked into the river Oulujoki (65.01°N, 25.27°E) draining into the Bothnian Bay. The broodstocks maintained at the LUKE hatcheries are renewed every few years with new broodstocks created from mature adults caught during their spawning migrations in their respective rivers. The parental salmon of our experimental cohorts were from first-generation hatchery broodstocks whose parents had successfully completed a marine migration.

To create our experimental cohorts, male and female salmon with homozygous *vgll3* genotypes (EE and LL) were crossed in multiple 2 x 2 factorials (13 and 17 factorials for the Oulu and Neva populations, respectively) with each factorial resulting in four families where all offspring within a family had the same *vgll3* genotype (EE, EL/LE or LL). For heterozygous EL and LE *vgll3* genotypes, the allele inherited from the mother is listed first, but in this study, the heterozygotes are treated as one genotype, EL.

The eggs were fertilized in October 2017 at the Viikki campus of the University of Helsinki (60.23°N, 25.02°E) and moved to the common-garden experimental facilities at Lammi Biological Station (LBS; 61.05°N, 25.04°E, Lammi, Finland) at the first-feeding stage in February -March 2018. Alevins were allocated to six circular tanks (Ø277 cm, maximum water volume 4.6 m^3^), with evenly mixing populations, *vgll3* genotypes, and families in each tank. Continuous water flow into the tanks was created by pumping water (UV-sterilized and heated by ∼1°C prior to entering the tanks) from the nearby lake Pääjärvi, with the water temperature in the experimental tanks following the lake’s natural temperature fluctuations (Figure 1). The average water temperature in the tanks during the year prior to spawning was 6.82 ± 3.17°C (min 2.14°C, max 18.10°C) and across the whole experiment 7.68 ± 3.93°C (min 0.22°C, max 18.69°C). In March 2021, heating of the incoming water was terminated, and individuals were redistributed into 12 tanks in order to reduce biomass. Light conditions in the experimental tanks followed the local photoperiod.

**Figure 1.**
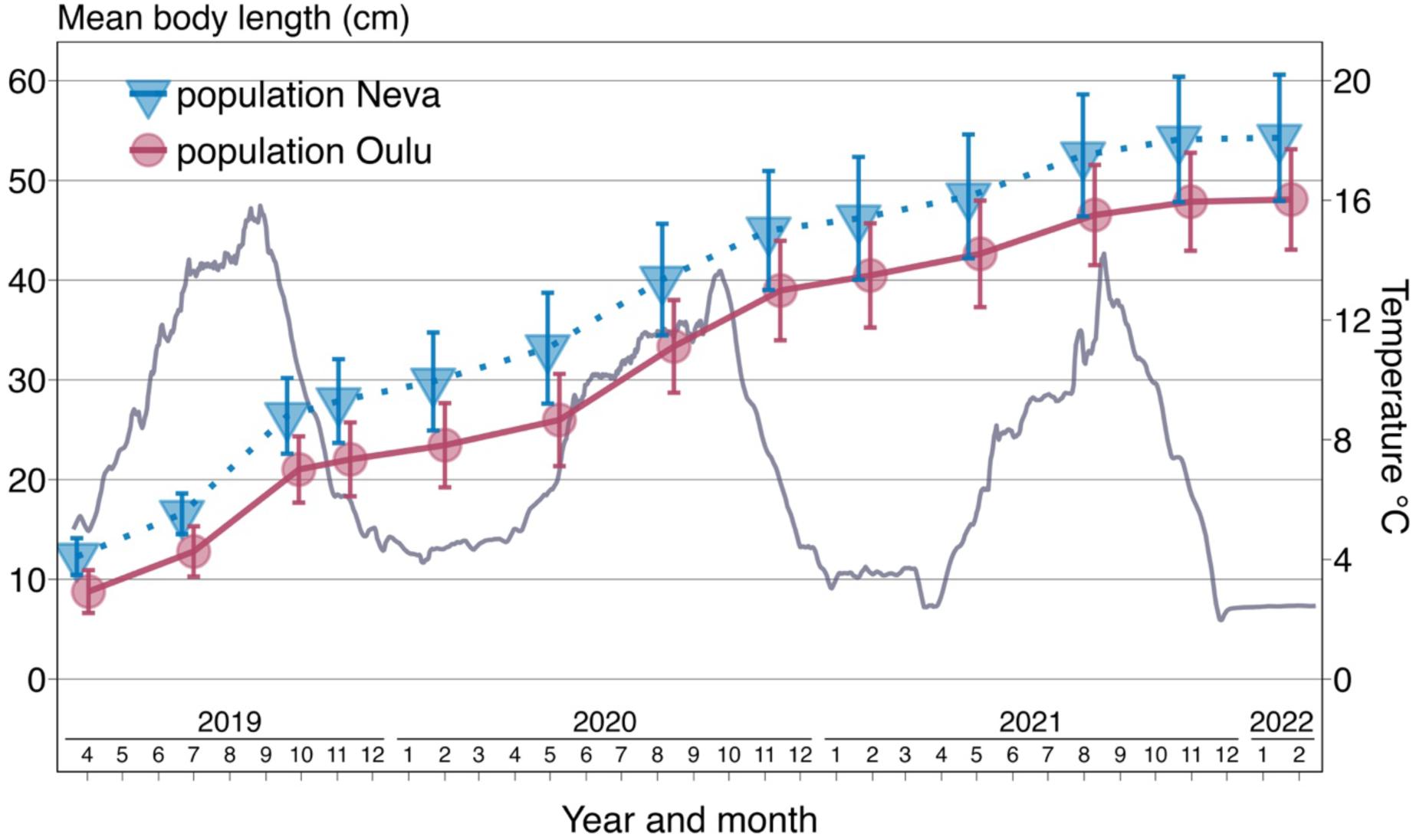
Growth of female Atlantic salmon represented as mean body length in centimetres during the common-garden experiment from April 2019 to February 2022. Body length is plotted separately for the Oulu (red circles) and Neva (blue triangles) populations, with feed treatments pooled together. Error bars indicate one standard deviation. The grey line represents the seasonal temperature fluctuations (7-day rolling average) in the experimental tanks.

### 2.2 Tagging, phenotypic measurements, and genetic analyses

In April 2019, all salmon were tagged with 12 mm passive integrated transponder (PIT) tags enabling identification of individuals and thus tracking of individual size and growth from that point onwards. Fin clips were taken for genetically determining the family, and thereby population and *vgll3* genotype of the individuals as described in Åsheim et al., (2023).

All fish were measured four times per year with approximately three-month intervals between each measurement. Prior to measuring, the salmon were anaesthetized with sodium bicarbonate-buffered tricane methanesulphonate (MS-222; concentration 0.125 g l^-1^). During the measurements, wet body mass (in g) and fork length (in mm) of the individuals were recorded. These body size measurements were then used to calculate Fulton’s condition factor (K) for all individuals with the formula K = 100 x (*mass (g)*/*fork length^3^ (cm)*) (Heincke, 1908).

### 2.3. Feed treatment

In July 2019, one year and four months after exogenous feeding had commenced, a feed treatment was started, whereby the salmon were fed either a control or a low-fat feed. The control feed was standard commercial fish feed (Hercules Baltic blend; Raisioaqua, Raisio, Finland), whereas the low-fat feed was the same standard feed, but manufactured with a lower fat content. This lower fat content resulted in ∼22% lower energy content (kJ/100 g) for the low-fat feed (feed content analyses for each feed batch were conducted by Synlab Oy, Karkkila, Finland). Feed was administered *ad libitum* to the tanks of both treatments using automatic feeders (Arvo-Tec Oy, Huutokoski, Finland) during daylight hours and the feed pellets were size-matched with the body size distribution of fish in each tank. By the spawning season of autumn 2021, the fish had been fed on one of the two diets for approximately two years and four months.

### 2.4 Maturation

Maturation status of the females (*n* = 753) was assessed in November 2021 when the salmon were four years old. During the previous 2020 spawning season, there were 59 mature females in our experimental cohorts (Maamela et al., 2023). The previously mature individuals that were still alive in November 2021 (*n* = 21) were removed from the analyses presented here due to their low number and the complications the inclusion of repeat spawning and previously mature individuals would have introduced in the analyses. Thus, none of the females included in this study had matured previously and this study is only assessing first-time maturation.

During the spawning of 2021, there were females from 46 families of the Oulu population and 39 families of the Neva population from 13 and 14 2 x 2 factorials, respectively. Maturation was assessed between 1^st^ – 18^th^ November 2021 and between 31^st^ January – 8^th^ February 2022 based on external characters as in Maamela et al., (2023). Briefly, females with an extended cloaca and releasing eggs when gentle pressure was applied to the abdomen were considered mature and females with an absence of these signs as immature. The majority of the mature females were releasing eggs in November. The February time point was used to confirm that the females that were maturing in November, but that were not yet releasing eggs, indeed matured later during the spawning season. Fecundity (egg number) was measured in a sub-sample of 318 females in November 2021. The sub-sampled individuals were randomly chosen from the mature females with the aim to have all *vgll3* genotypes represented as equally as possible within feed treatments for both populations. Since our experiment included more fish from the Oulu population (*n* = 469) compared to the Neva population (*n* = 284), the study has a higher representation of Oulu individuals for the fecundity analysis. In total, we analysed fecundity for 248 and 74 females from the Oulu and Neva populations, respectively. The number of sampled females per *vgll3* genotype, feed treatment, and population are summarised in Table S1.

The females were hand stripped by gently massaging the abdomen to extract the eggs. The total wet mass (g) of the egg clutch was obtained by weighing after straining off the ovarian fluid with a metal sieve (mesh size 2 mm). Three replicates of 10 eggs each were weighed (to 0.0001 g accuracy) to obtain individual egg wet weight. Fecundity of the individuals was calculated by dividing the total egg clutch mass with the mean individual egg wet weight.

### 2.5. Statistical methods

#### 2.5.1 Samples used in the models

A total of 730 female salmon which were spawning for the first time, and for which phenotypic, genotypic, and population information were available, were included in the final models for maturation probability, body size, and body condition. An overview of the number of female salmon in the different treatments are summarized together with the observed maturation rates in Table S2.

#### 2.5.2 Common model structures

We analysed the effect of diet, *vgll3*, and population on maturation, body condition, body length, and fecundity with generalized and general linear mixed effect models. The common fixed variables used in all models were feed treatment (control, low-fat) and population (Neva, Oulu), both coded as two-level factor variables. To test for additive effects of *vgll3* genotypes, the genotypes were coded as -1, 0, and 1 for the LL, EL, and EE genotypes, respectively. Though dominance effects at the *vgll3* locus on age at maturity have been documented previously in the wild (Barson et al., 2015; Czorlich et al., 2018; Mobley et al., 2024), none have been detected in controlled conditions for males (Åsheim et al., 2023; Debes et al., 2021). Similarly, for female salmon, the previously found dominance effects have been weaker or population-or age-specific compared to males (Barson et al., 2015; Czorlich et al., 2018) and in line with this, other studies have shown the effect of *vgll3* to be completely additive in females (Miettinen et al., 2024; Raunsgard et al., 2023). Therefore, we excluded dominance effects in our models in order to reduce model complexity and aid model fitting. Body length and body condition were mean-centered and standard deviation scaled before being included in the models. We used body size and body condition measurements taken in May 2021, approximately five months prior to spawning, in all analyses. This spring timepoint was chosen for the measurements as we wanted to model the effect of available energy for reproduction before the individuals had allocated energy towards gonad development for the coming spawning season (Åsheim et al., 2023; Debes et al., 2021).

#### 2.5.3 Maturation model

Maturation probability was analysed using a generalized linear mixed effect model with maturation coded as a Bernoulli-distributed binary variable (0 for immature, 1 for mature) with a logit-link function. The *vgll3* genotype, population, feed treatment, body length, and body condition were included as fixed effects in the model. In addition, we included two-way interactions between *vgll3* genotype, population, and feed treatment to assess genotype-by-environment (GxE) effects on maturation probability. An interaction between *vgll3* and body condition was also included as previous studies have found *vgll3* to be associated with the body condition of Atlantic salmon (Debes et al., 2021; House et al., 2023). Initially, we also included a three-way interaction between *vgll3*, population, and feed as the observed maturation rates indicated that such an interaction might be present (see Figure 2). There was no clear three-way interaction effect on maturation (the 95% credible interval overlapped with zero) and we removed this interactive effect from the final model in order to aid model fitting.

**Figure 2.**
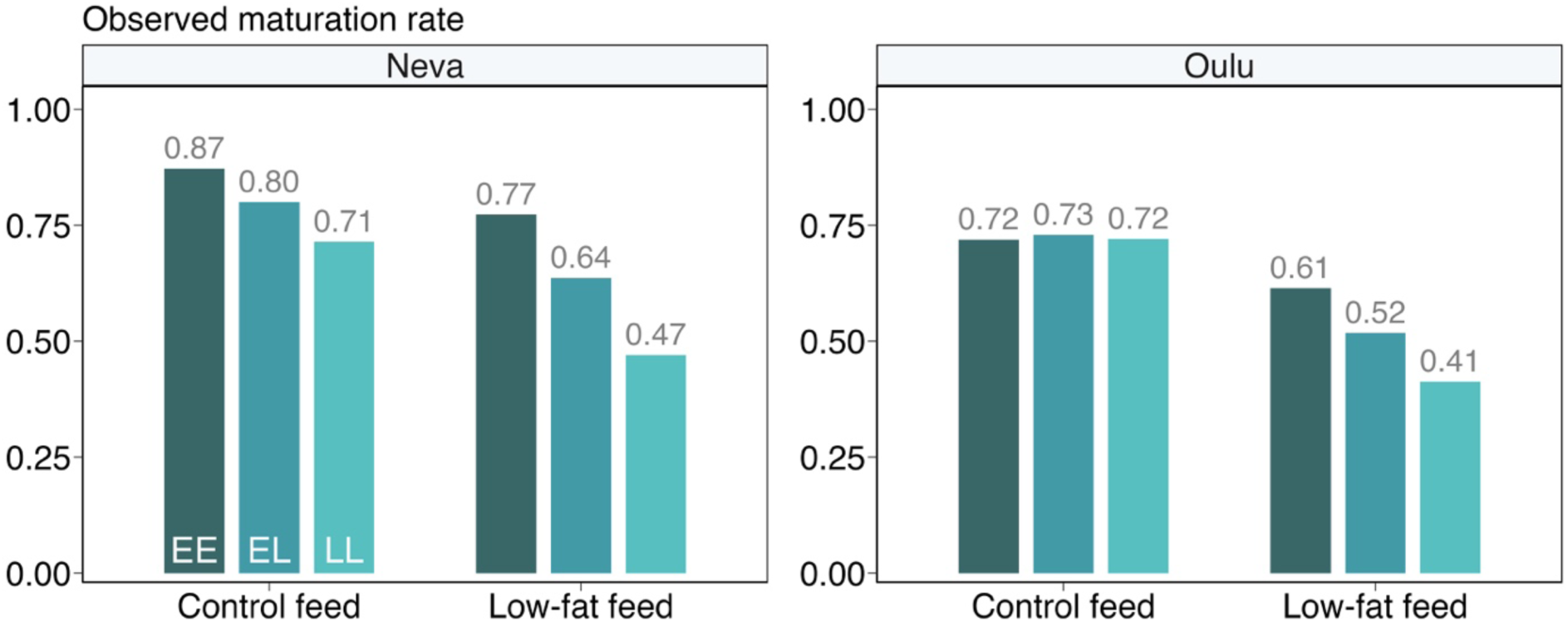
Observed proportions of *vgll3* genotype-specific maturation in female Atlantic salmon from two populations (Neva, Oulu) in two feed treatments. The colours indicate the three different *vgll3* genotypes (dark blue = EE, blue = EL, light teal = LL). The exact maturation rates and number of salmon per group are presented in Table S2.

Family structure and relatedness among individuals was accounted for using the animal model approach (Wilson et al., 2010) by including the effect of an individual animal in the model as random effect. This effect of an individual was fitted using the additive genetic relatedness matrix obtained from the pedigree. Our pedigree included family relationships up to the grandparental level. These grandparental individuals were the Atlantic salmon that successfully completed a sea migration and were the founders of the broodstock created by LUKE (see information about broodstocks above and in Åsheim et al., 2023). By utilizing the pedigree-based relatedness matrix to correlate the additive genetic random effects in the model, we were able to obtain estimates of additive genetic variance (V_A_) which were then used to calculate heritability of maturation. Heritability (*h^2^*) was calculated by dividing the sum of V_A_ and additive variance contributed by *vgll3* (V_A,*vgll3*_) by the total phenotypic variance estimate (V_P_) calculated as the sum of the different variance components (V_P_ = V_A_ + V_A,*vgll3*_ + V_tank_ + V_residual_). V_A,*vgll3*_ was estimated as in Debes et al. (2021) and in Åsheim et al., (2023) using the formula V_A,*vgll3*_ = 2*pq*(α*_vgll3_*)^2^, where *p* and *q* are the *vgll3* allele frequencies in the data and α*_vgll3_* is the weighted average of the *vgll3* model regression coefficient estimates within treatment groups (i.e., populations and feed treatments). V_tank_ was obtained from the model random tank effects and V_residual_ was fixed to π^2/3^. V_A,*vgll3*_ was also used to calculate SNP heritability (*vgll3*’s contribution to total phenotypic variance) by dividing V_A,*vgll3*_ by V_P_.

Raw parameter estimates from the maturation model were converted from log odds ratios to odds ratios (*e*^log odds ratio^) in the text, thus giving the relative estimate of change in the odds of maturing. Odds are the probability of maturing divided by the probability of not maturing, and ranges from zero to infinity, which is simpler to model compared to probability which is bound between zero and one.

#### 2.5.4 Body condition, length, and fecundity models

We used linear mixed effect models with an identity-link function to investigate the effects of the focal traits and treatments (*vgll3*, population, feed treatment) on body condition, body length, and fecundity. Similar to the maturation probability model, we added two-way interactions between all fixed effects.

In the fecundity model, we included body length and body condition of the individuals for which we had fecundity data as additional covariates as these traits are known to influence egg number in fish (Barneche et al., 2018). Based on the results of the body condition and length models (see Results), we also included two-way interactions between population and body length as well as population and body condition. An interaction between *vgll*3 and body condition was also added. Fecundity was mean-centered and standard deviation scaled before being included in the model to aid model fitting. In the plotted results, predicted fecundities have been back-transformed to the original scale.

#### 2.5.5 Modelling technicalities and assessment of model fits

All models were fit using Bayesian statistical methods. The models used four Hamiltonian Monte Carlo chains each run for 3,000 iterations with the first 500 iterations discarded as warm-up. Thus, the models resulted in 10,000 realisations of the posterior distribution for each response variable. For all models (maturation, fecundity, body condition, and body length), we set all priors for the intercept, parameter estimates, and random factors as fairly non-informative normal distributions with a mean of zero and standard deviation of one. These priors assume a 50% probability of maturation at the intercept and that there is no effect of any of the variables included in the model.

Models were assessed for convergence and autocorrelation based on a visual inspection of the trace and autocorrelation plots and by using R-hat values. All R-hat values were below 1.05 and the plots showed proper mixing of the chains. No autocorrelation was detected. In addition, model fits were checked by calculating pareto k diagnostics and performing a visual posterior predictive check. The pareto k diagnostics found no highly influential points for the maturation, body condition or body length models, but one influential point (0.3%, k > 0.7) was found in the fecundity model.

#### 2.5.6 R packages used

All data analyses were done using Rstudio version 2023.12.1 running R v4.4.0 (R Core Team, 2024). Data management and visualisation was done using the *tidyverse* package v2.0.0 (Wickham et al., 2019). The *loo* package v2.7.0 (Vehtari et al., 2017, 2024; Yao et al., 2018) was used to calculate pareto k model diagnostics. All Bayesian models were run using *rstan* v2.19.3 (Stan Development Team, 2020) via the *brms* package v2.17.0 (Bürkner, 2017, 2018, 2021).

### 2.8. Ethical statement

The experiments were approved by the Project Authorisation Board (ELLA) on behalf of the Regional Administrative Agency for Southern Finland (ESAVI) under experimental licence ESAVI/2778/2018.

## 3.0 Results

### 3.1 Overall maturation rates

Overall, 66.8% of the 730 female salmon in our experiment were mature with population-specific maturation rates being 74.6% for Neva and 62.3% for Oulu individuals, respectively (Figure 2). Across populations, the observed *vgll3* genotype specific maturation rates were 72.9%, 65.3%, and 59.3% for the vgll3 EE, EL, and LL genotypes, respectively. Maturation rates were lower in the low-fat feed treatment (59.0%) compared to 75.7% in the control feed treatment. The highest maturation rate (87.2%) was observed for the *vgll*3*EE individuals of the Neva population that were fed on the control diet. This was more than double the lowest maturation rate of 41.3% for *vgll3*\*LL Oulu individuals from the low-fat feed treatment (Figure 2, Table S2).

### 3.2. Statistical results for maturation, body length, and body condition models

#### 3.2.1 *vgll3* genotype effects

*Vgll3* genotype influenced maturation probability of female salmon (Figures S1, 2 and 3, Table S5). The effect of *vgll3* was additive with each added E allele increasing the relative odds of maturation by 2.41 fold ([95% CI: 1.07, 5.42], for Neva individuals in the control-feed group, of average body condition). We did not detect clear interactions between *vgll3* and feed or between *vgll3* and body condition on the probability of maturation (Figure 3). There was, however, an uncertain interaction (the 95% CI overlapped with zero, with 97.1% of the total posterior distribution being below zero and 2.9% above zero) between *vgll3* and population, whereby the additive effect of *vgll3* on maturation was lower for the Oulu individuals and their predicted maturation probability lacked any *vgll3* effect (Figure 3). This interaction was likely driven by the Oulu individuals in the control feed treatment where the effect of *vgll3* genotype on maturation was not evident (Figures 2 and 3).

**Figure 3.**
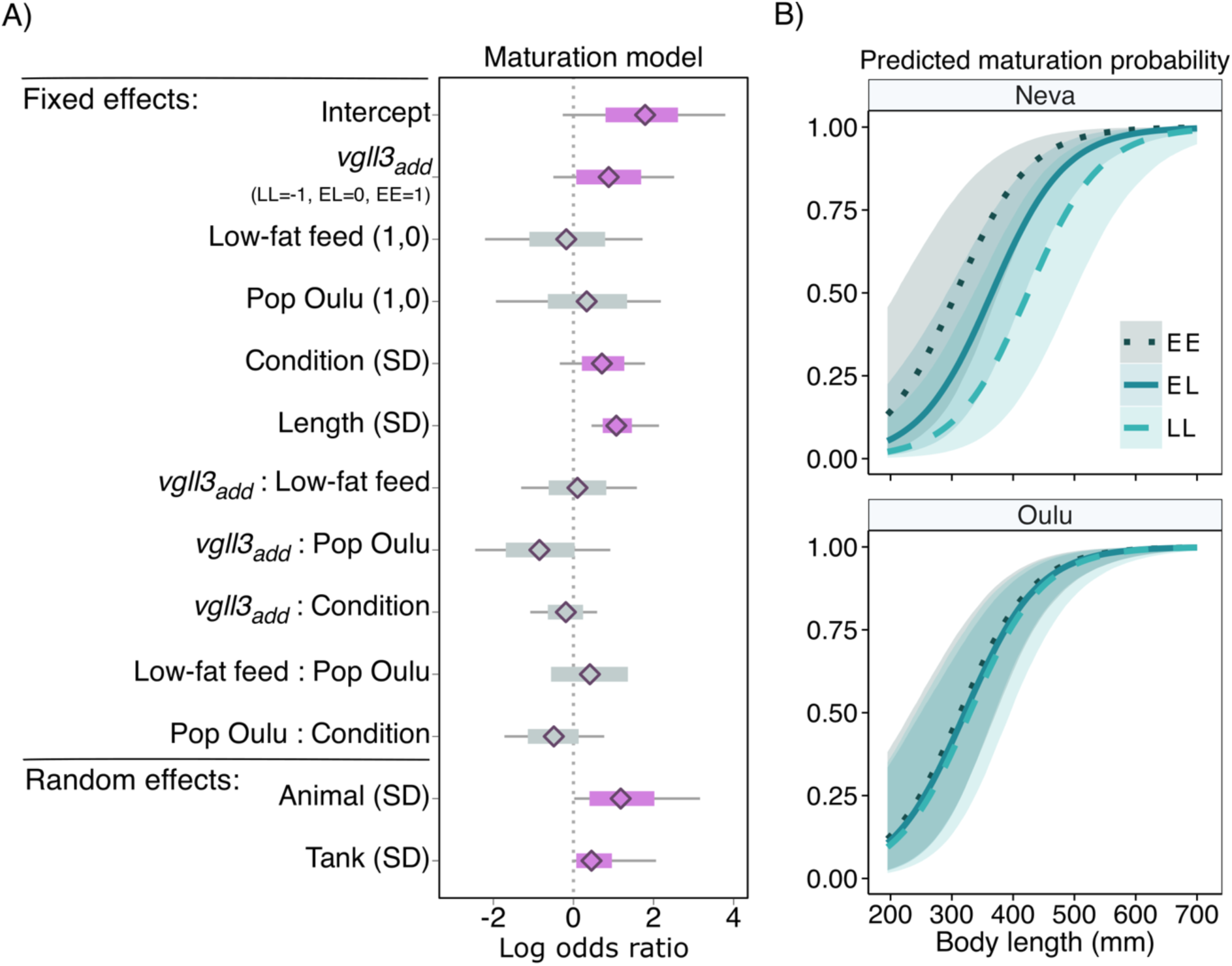
A) Parameter estimates (log odds ratio) and 95% credible intervals (CI) for the fixed and random effects obtained from the maturation model. The diamonds indicate the mean parameter estimates calculated from the posterior samples, the thick bars indicate the 95% CIs, and the thin bars the 100% CI. The thick bars are coloured purple if the 95% CI does not overlap zero. The scales of the fixed and random effect model parameters are indicated in the parentheses after the variable name. For categorical variables, the levels are 0 = Neva and 1= Oulu for population, and 0 = control and 1 = low-fat for feed. Continuous variables were centered and standard deviation (SD) scaled which leads to model estimates indicating the effect of increasing or decreasing the variable by one SD. Model intercept was set to 0 for all variables. Random effects standard deviations indicate the magnitude of variation among tanks and breeding values. Full model results are presented in table S5. B) Predicted maturation rates for female Atlantic salmon with different *vgll3* genotypes (dotted dark blue = EE; solid blue = EL; dashed teal = LL) from two different populations (Neva; Oulu) along body length (mm). Predictions were estimated using the maturation model with feed treatment set as the low-fat feed and body condition as the mean of all the female salmon included in the analysis. The shaded areas along the prediction lines represent the 95% CI.

V*gll3* genotype did not influence body length or body condition in the spring prior to spawning (Figures S3 and S4, Tables S6 and S7).

#### 3.2.2 Feed

Diet did not have a direct effect, meaning an effect which is independent of the effect of body condition, on maturation probability for females. The model estimate for the effect of feed on maturation was close to zero with a 95% credible interval that included zero (Figure 3). Diet did affect body condition, with the low-fat diet being associated with lower body condition (model estimate: -0.88 [95% CI: -1.20, -0.54]; Figure S3). There was an interaction between feed treatment and population on body condition whereby the decrease in body condition due to the low-fat diet was slightly larger in Oulu than in Neva individuals (model estimate: -0.22 [95% CI: -0.44, -0.01]; Figure 4). No effect of feed was found on either body length (Figure S4) or fecundity (Figure 4).

**Figure 4.**
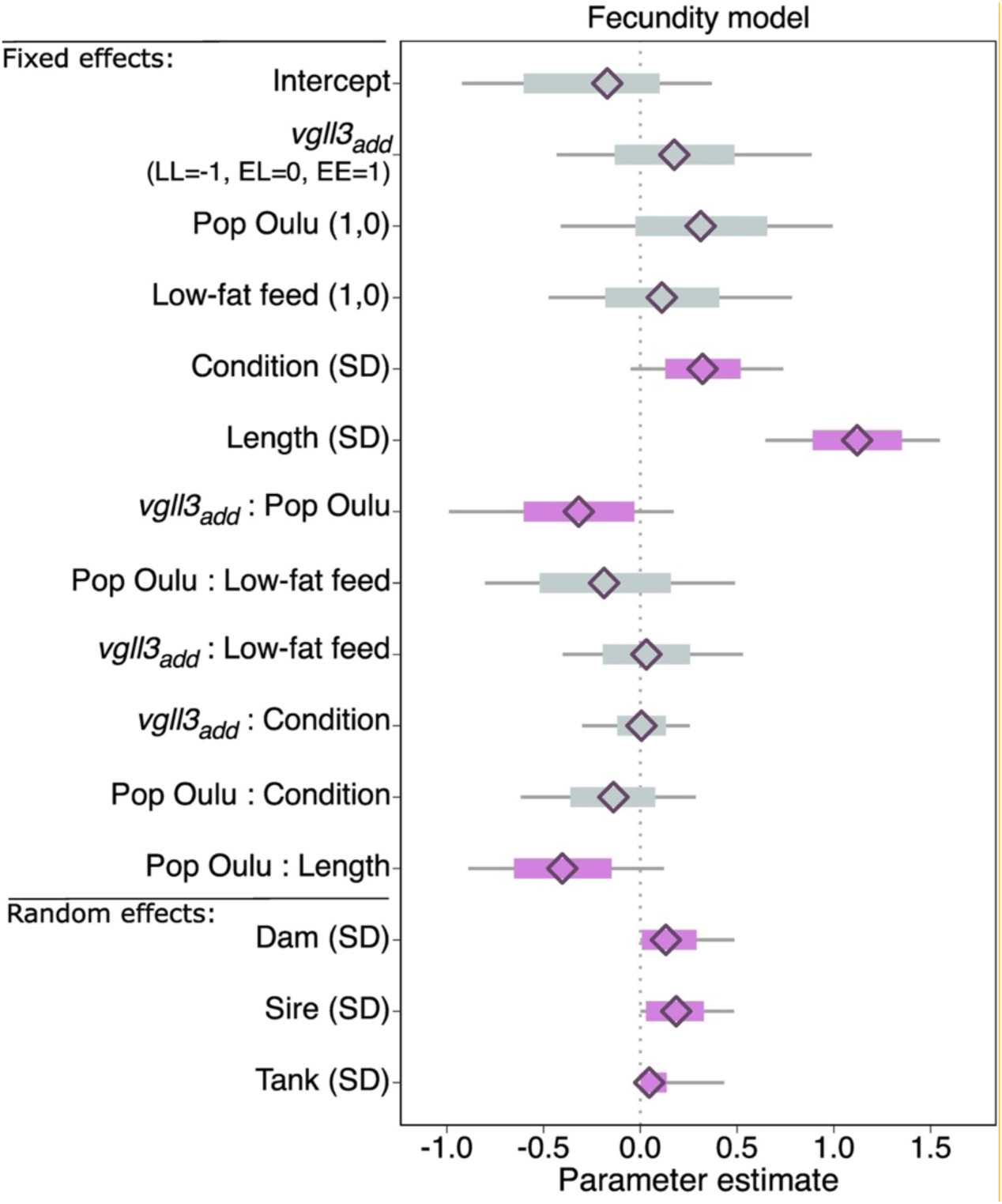
Parameter estimates and 95% credible intervals for the fixed and random effects obtained from the fecundity model. The scales of the fixed and random effect model parameters are indicated in the parentheses after the variable name. For categorical variables, the levels are 0 = Neva and 1= Oulu for population, and 0 = control and 1 = low-fat for feed. Continuous variables were centered and standard deviation (SD) scaled which leads to model estimates indicating the effect of increasing or decreasing the variable by one SD. Model intercept was set to 0 for all variables. Random effects standard deviations indicate the magnitude of variation between tanks and between parents. The diamonds indicate the mean parameter estimates calculated from the posterior samples, the thick bars indicate the 95% CIs, and the thin bars the 100% CI. The thick bars are coloured purple if the 95% CI does not overlap zero.

#### 3.2.3 Population

We did not find a direct effect of population on maturation probability (Figure 3), but population influenced both body length (model estimate: -1.00 [95% CI: -1.35, -0.65]; Figure 1, Table S7) and body condition (model estimate: -0.56 [95% CI: -0.87, -0.25], Figure S3, Table S6). Oulu individuals were, on average, smaller (Figure 1) and had lower body condition (Figure S7) than the Neva individuals. For example, in May 2021 the mean body length of Oulu individuals was 42.6 ± 0.25 cm compared to 48.4 ± 0.38 cm for Neva (Table S3).

#### 3.2.4 Effects of body size and body condition on maturation

Both larger body size and higher body condition in the spring prior to spawning were associated with higher probability of maturation (Figures 3 and S2). Body length had a slightly larger influence on maturation compared to body condition. An increase of one standard deviation in body length increased the odds of maturation by 2.92 fold [95% CI: 2.08, 4.30] whereas the effect of one standard deviation in body condition increased maturation odds by 2.04 fold [95% CI: 1.23, 3.55].

#### 3.2.5 Heritability

Heritability (*h^2^*) of female maturation was 0.295 [95% CI: 0.064, 0.543]. We estimated the contribution of *vgll3* to phenotypic variance to be 0.02 [95% CI: 0.00, 0.065].

### 3.3 Fecundity

Fecundity ranged from 728 to 6937 eggs with the average being 2910 (Figure 5, Table S4). As expected, body size affected fecundity with longer body length resulting in higher egg number (model estimate: 1.12 [95% CI: 0.9, 1.35], Figures 4 and 5, Table S8). Population had an influence on the number of eggs a salmon produced; the mean fecundity of a Neva individual was 3750 whereas mean fecundity was 2670 for an Oulu individual (Table S4). Interestingly, the fecundity-increasing effect of body length was not as strong for the Oulu individuals compared to the Neva as indicated by the interaction term between population and length (model estimate: -0.40 [95% CI: -0.65, -0.15], Figure 4, Table S8). In addition, there was an interaction between population and *vgll3* genotype, where the additive effect of *vgll3* on fecundity was not as strong for the Oulu compared to Neva individuals (model estimate: -0.32 [95% CI: -0.61, -0.04], Figures 4 and S5, Table S8).

**Figure 5.**
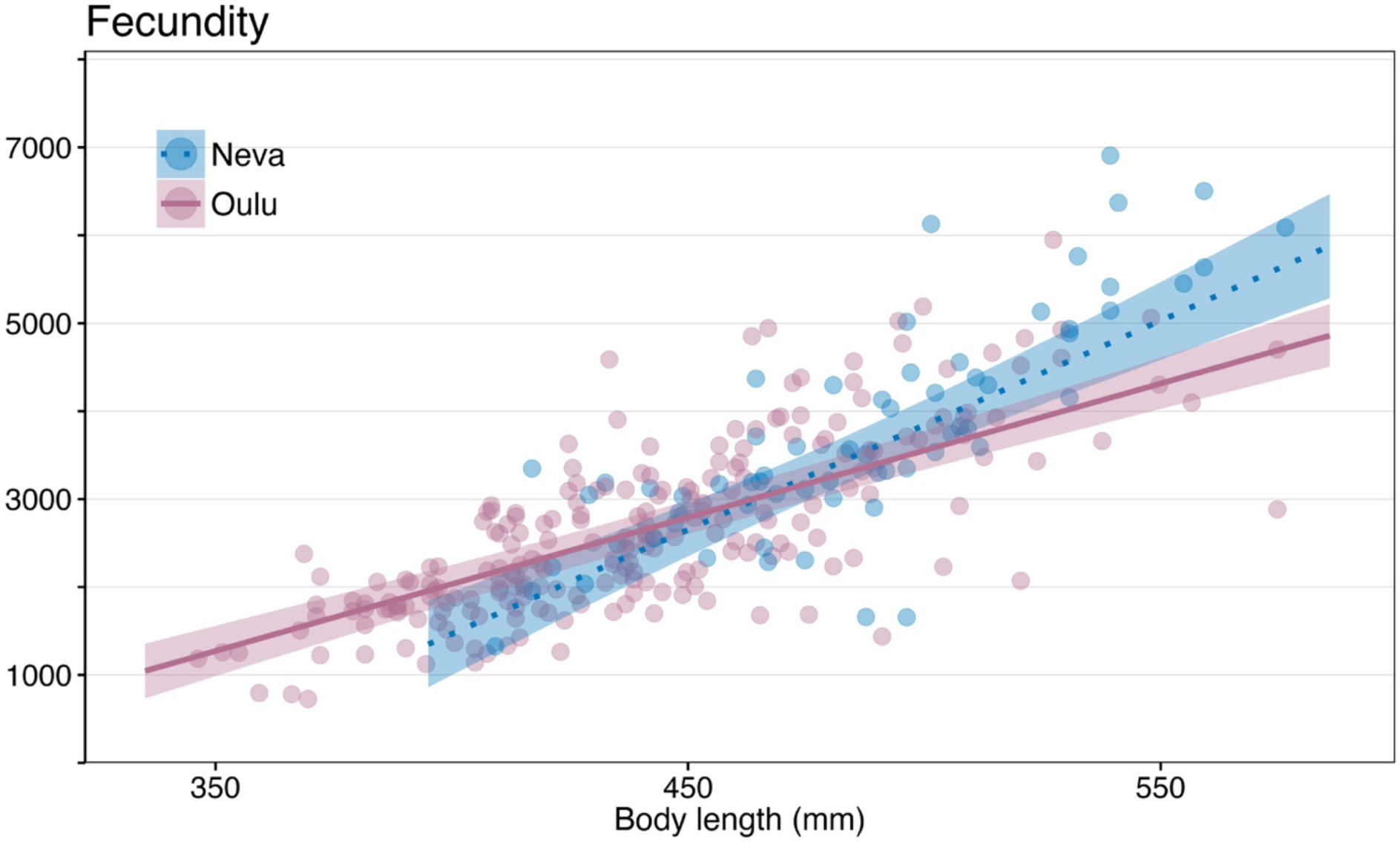
Fecundity (number of eggs) per mature individual plotted against body length in the spring prior to spawning. The points are the observed fecundity coloured by population (blue = Neva, red = Oulu). The trendlines indicate the predicted fecundity calculated using the fecundity model. For the predictions, *vgll3* genotype was set as EL and feed as low-fat feed. Body condition was set as the mean of all the female fish included in the fecundity model. The shaded areas around the trendlines are the 95% credible intervals.

## 4.0 Discussion

Female Atlantic salmon have seldom been included in common-garden experiments studying maturation, which has left a gap in our knowledge of the genetic and environmental factors influencing age at maturity. In this study, we reared female salmon until four-years-old in a common-garden experimental setting where we manipulated the energy content of their diets and assessed the probability to mature for the first time at this age. We found genetic effects on maturation and maturation-related traits both at a single-locus and at a population-of-origin level. *Vgll3* genotype influenced maturation probability, whereas population had an indirect effect on maturation via body condition, and an effect on fecundity. Additionally, higher energy content diet was associated with higher body condition. As higher body condition was also associated with higher probability of maturation, diet had an indirect effect on maturation via the available energy resources that could be allocated to reproduction.

### 4.1 *Vgll3* and female maturation

We demonstrate that the *vgll3* locus influences female Atlantic salmon maturation in a controlled experimental setting similar to how it affects female maturation in the wild (e.g., Barson et al., 2015; Besnier et al., 2024; Czorlich et al., 2018; Miettinen et al., 2024; Raunsgard et al., 2023). The effect of *vgll3* was additive with each additional copy of the *vgll3*\*E allele increasing the probability of maturation. Overall, the *vgll3* effect on maturation was very similar across feed treatments and populations, except for Oulu individuals in the control feed treatment where no apparent effect of *vgll3* was found; the observed maturation rate in this group was approximately 72% regardless of *vgll3* genotype and there was no difference in predicted maturation probabilities between genotypes for the Oulu individuals. This relatively high maturation probability and lack of association with *vgll3* could indicate a population-specific maturation threshold linked with body condition. On average, the Oulu individuals had lower body condition compared to Neva. If the Oulu salmon have a lower body condition threshold for maturation compared to Neva salmon, then our result could indicate that when feed energy content is sufficiently high and the individuals are able to acquire abundant energy reserves, the effect of *vgll3* on maturation is bypassed by other environmental and genetic mechanisms mediating maturation (Besnier et al., 2024). In a previous study, *vgll3* was not associated with female age at maturity in a commercial Mowi aquaculture strain salmon in a common-garden experiment (Ayllon et al., 2019). In that study too, the authors hypothesized that excess availability of feed and other environmental conditions under farming conditions and the strain’s long history of domestication, which all influence growth rate, may have overridden the potential effect of *vgll3* on maturation.

To better understand the genetic basis and adaptive potential of age at maturity in female Atlantic salmon, we calculated both heritability and *vgll3*’s contribution to phenotypic variance of maturation. Heritability was estimated to be approximately 0.295 in our study. This estimated heritability for females was lower compared to heritability of first-time maturation for the male salmon (estimated to be 0.68) in our experimental cohorts (Åsheim et al., 2023), but in line with the median estimate of 0.21 calculated for multiple salmonid species for both sexes (Carlson & Seamons, 2008). Overall, our estimate of maturation heritability suggests a moderate response to selection. In addition to heritability, we calculated *vgll3*’s contribution to phenotypic variance of first-time maturation and found the contribution to be only ∼2%. In contrast, the contribution of *vgll3* on phenotypic variance in males from the same common-garden experiment was ∼13% (Åsheim et al., 2023). Both the lower heritability of maturation and *vgll3*’s contribution to it for females indicates that the additive genetic component of age at maturity is not as prominent in female compared to male Atlantic salmon. This smaller additive genetic contribution highlights the importance of environmental factors on female maturation in our experiment, though we cannot exclude the potential influence of non-additive genetic factors (e.g., epistasis) on maturation (Roff & Emerson, 2006). Small sample sizes prevented us from fitting animal models that could have quantified the effect, if any, of epistatic effects. The low heritability of maturation could be linked to age at maturity being an important life-history trait as traits connected with fitness often exhibit lower heritabilities compared to morphological traits (Falconer & Mackay, 1996; Mousseau & Roff, 1987). In Atlantic salmon, both sexes get a reproductive fitness benefit from older age at maturity (Mobley et al., 2020, 2024), and the mean sea age at maturity for the sexes can differ between populations due to local adaptation (Erkinaro et al., 2019; Persson et al., 2023; Tréhin et al., 2024). On average, age at maturity is likely to be higher for female salmon compared to males partly due to female reproductive success being more tightly linked with body size and, thus, with age (Czorlich et al., 2018; Erkinaro et al., 2019; Fleming et al., 1996; Fleming & Einum, 2011; Mobley et al., 2020). Interestingly, differences between the sexes in how *vgll3* influences age at maturity, where the effect of *vgll3* on age at maturity appeared to be smaller for females than for males, have also been reported for a wild Norwegian Atlantic salmon and a Baltic salmon population (Besnier et al., 2024; Miettinen et al., 2024). Moreover, in their study comparing historical and contemporary salmon samples from river Etne in Norway, Besnier et al., (2024) noted that the influence of *vgll3* on age at maturity was almost completely absent in contemporary samples from the 2010s, while an effect was still present in the samples from the 1980s. Age at maturity is mediated by multiple loci (Sinclair-Waters et al., 2020). With this polygenic nature of the trait in mind together with the findings of the sex-specific patterns in *vgll3*’s effect on maturation in our study and in the studies in the wild populations (Besnier et al., 2024; Miettinen et al., 2024), age at maturity in female Atlantic salmon might have a larger environmental component compared to males.

### 4.2 Other factors influencing maturation

In many taxa, individuals require sufficient energy stores for initiating the maturation process both in terms of metabolic costs and direct energetic costs allocated to the offspring (Ginther et al., 2024). Age at maturity in Atlantic salmon is often thought to be, and is modelled as, a threshold trait, where the initiation of maturation process is triggered by an underlying (continuous) liability trait, such as growth rate, body size or somatic energy reserves, surpassing a genetically predetermined threshold value (Hutchings, 2021; Lynch & Walsh, 1998; Piché et al., 2008; Thorpe et al., 1998). Due to the large reproductive fitness advantage they receive from delaying maturation, female Atlantic salmon have a higher growth and body size threshold for maturation compared to males (Hutchings & Jones, 1998; Jonsson & Jonsson, 2011; Tréhin et al., 2021). As expected, we found both larger body size and higher body condition to be associated with higher probability of maturation.

Though an effect of *vgll3* on body condition has been demonstrated previously in males (Debes et al., 2021; House et al., 2023), we were unable to find an association in this study in either the maturation or the body condition model. As shown by House et al., (2023) and Debes et al., (2021), body condition is not a static trait but rather changes throughout the year and the magnitude of differences in body condition between the *vgll3* genotypes also changes depending on sampling timepoint. We chose to measure body condition in May as condition in spring and early summer is a predictor of maturation status in autumn for salmon (Kadri et al., 1997). Moreover, this is the time point where an association between *vgll3* and body condition has been previously found in males (Debes et al., 2021), though we cannot exclude the possibility that *vgll3* may influence body condition differently in females than in males.

We did not find an independent effect of feed treatment on maturation probability, although it has earlier been shown that diet can influence age at maturity in Atlantic salmon (Jonsson et al., 2013). We did, however, detect a possible indirect feed effect via body condition. The individuals that had been fed the lower energy low-fat diet for approximately 2.3 years before spawning had lower body condition compared to the individuals receiving the standard, higher energy diet. The lower body condition in the low-fat diet treatment suggests that immature individuals in this treatment were unable to acquire sufficient energy stores for maturation due to reduced feed quality causing them to be unable to reach the body condition threshold needed for maturation, thus delaying their maturation. The effect of diet was only found on body condition and not on body length. The lack of diet effect on body length indicates that the salmon in the higher energy feed treatment were allocating their acquired energy to energy storage rather than to growth.

We found that population of origin influenced body length and body condition. Individuals from the Oulu population not only had shorter body length, but they also had lower body condition compared to Neva individuals. In addition, there was an interaction between diet and population whereby Oulu individuals displayed a larger reduction in body condition in the low-fat treatment than Neva individuals. As individuals from both populations were grown in the same environmental conditions and mixed evenly between tanks and feed treatments, the observed differences indicate that there are possible genetic or epigenetic differences between the populations in their way of utilizing dietary energy resources.

### 4.3 Fecundity

The positive correlation between female body size and fecundity is well established in various fish species, including Atlantic salmon (Barneche et al., 2018; Fleming, 1996; Hanson et al., 2020; Moffett et al., 2006; Thorpe et al., 1984). The results from this study also demonstrated this relationship between individual body size and egg number with larger females producing more eggs. Additionally, we found that body condition in the spring prior to spawning also influenced fecundity. The individuals that had a higher body condition in the spring also produced more eggs. Both body size and body condition were found to affect fecundity in the same model, which indicates that both traits may be important for fecundity and that the mechanisms of how they influence egg number can be different. For example, body condition can, alternatively or in addition to representing energy resources, reflect differences in body shape and thus, space available for eggs.

Oulu individuals produced fewer eggs for a given body length compared to the Neva individuals. Differences in fecundities between populations, after accounting for body size or age, have been reported for wild salmon in Europe (de Eyto et al., 2015; Fleming, 1996; Hanson et al., 2020; Klemetsen et al., 2003). This variation between rivers could arise due to the spawning history of the female (e.g., first-time vs. repeat spawner) affecting egg size and/or number (Reid & Chaput, 2012) or due to environmental variation either in the freshwater or marine environment utilized by the salmon (Jacobson et al., 2021). The finding of differing fecundities between the two populations in our study indicates that beyond environmental and age-specific effects, genetic factors between populations may influence the number of eggs an individual produces. However, we did not find a clear effect of *vgll3* on fecundity, as estimated by egg number, but there was a weak interaction between population and *vgll3* on fecundity — the additive effect of *vgll3* on fecundity was weaker for Oulu compared to Neva individuals. When interpreting these *vgll3* results from the model, it should be noted that there were fewer Neva individuals in our study cohort compared to Oulu individuals and in particular, fewer LL individuals from the Neva population (11 compared to 65 from the Oulu population). This could lead to an apparent effect of *vgll3* on fecundity when, in fact, it is just a statistical artifact arising from unbalanced experimental design. The lack of similar pattern in the observed fecundities suggests that this may be the case (Figure S4). We recommend follow-up studies to either confirm or reject this interactive effect.

### 4.4 The importance of studying female Atlantic salmon reproduction

In this study, we investigated genetic and environmental influences on maturation in two hatchery populations of female Atlantic salmon. We were able to validate the effect of *vgll3* on female maturation in controlled rearing conditions, an effect which so far has only been observed in the wild. Female salmon typically mature at an older age and larger size compared to males (Czorlich et al., 2018; Mobley et al., 2020) and thus, they have often been neglected in laboratory studies of maturation due to the financial, logistical, and labour costs associated with rearing the fish until maturation. The focus on male maturation may also partly be due to Atlantic salmon’s major role in aquaculture. There, early maturation is a larger problem in male compared to female salmon where maturing individuals shift their energy allocation from somatic growth to gonadal development which leads to smaller body sizes and financial losses (Taranger et al., 2010). Inadvertently, the exclusion of female salmon from long-term studies of maturation in controlled settings has led to incomplete knowledge of the processes underlying age at maturity. This is despite female age at maturity being a core component of many population management models. Exploitation of many wild Atlantic salmon stocks is managed by stock-specific spawning targets based on biological reference points where these targets are expected to be set at levels that prevent over-exploitation (Chaput et al., 1998; White et al., 2023). One important reference point used for the spawning targets is the number of eggs individuals in the managed population produce (Forseth et al., 2013; O’Connell & Dempson, 1995). Old and large females with high fecundity are known to have a large influence on population productivity across fish species (Hixon et al., 2014; Marshall et al., 2021). This connection between age at maturity and fecundity highlights the importance of female salmon for management and conservation, and of understanding how different factors influence age at maturity and maturation-related traits in female Atlantic salmon. In conclusion, the results of our study have broadened our understanding of the mechanisms behind Atlantic salmon age at maturity and we demonstrate that gene-by-environment interactions shape maturation and reproductive traits.

## Data availability statement

All data and code used in the analysis will be made available in Zenodo upon acceptance of the article.

## Benefit sharing statement

Benefits Generated: Benefits from this research accrue from the sharing of our data and results on public databases as described above.

## Author contributions

Conceptualization: CRP, PVD, KSM, AHH, ERÅ, KBM

Data curation: ERÅ, KSM

Formal analysis: KSM, RJO’S, ERÅ

Funding acquisition: CRP, KSM

Investigation: KSM, ERÅ, PTN, JMP, PL, KBM

Methodology: PVD, AHH, KSM, ERÅ, CRP, KBM

Project administration: CRP Resources: CRP, JE

Supervision: CRP, KBM

Visualization: KSM

Writing-original draft: KSM

Writing-review & editing: KSM, CRP, KBM, ERÅ, RJO’S, JMP, PTN, JE, PVD, AHH

## Acknowledgements

We wish to thank all the people who helped during this multi-year study. These include Jacqueline Moustakas-Verho, Ksenia Zueva, Marion Sinclair-Waters, Nico Lorenzen, Victoria Pritchard, Yann Czorlich, Spiros Papakostas, and Suvi Ikonen for their help with salmon gamete collection and performing the crosses that created the salmon used in this study, Natural Resources Institute Finland (LUKE) for access to the broodstocks, and staff of the Taivalkoski and Laukaa hatcheries for their help with gamete collection. Thank you to Nikolai Piavchenko and Noora Parre who cared for the fish at the Viikki campus. Susanna Airaksinen and Thomas Ginström and others at Raisioaqua Ltd are thanked for producing the low-fat feed used in the study. Suvi Ikonen, Andrés Salgado, Anna Toikkanen, Dorian Jagusch, Fin Morrison, Ike van Gestel, Jeferson Delgado Florez, Maël Le Gouellec, Markus Lauha, Mikko Immonen, Paul Bangura, Juho Kökkö, Matias Boehm, Simon Lecarte, Teemu Mäkinen, and Tiemen Jansen are thanked for their assistance with fish husbandry and sampling and John Loehr for assistance with logistics at the Lammi Biological Station. We thank Annukka Ruokolainen, Seija Tillanen, Shadi Jansouz, Iikki Donner, Morgane Frapin, and Tutku Aykanat for assistance with SNP genotyping and sex determination and Patrick Heidbreder, Nora Bergman, Xindi Huang, Amaïa Lamarins, and Morgane Frapin for useful discussions.

## Supplementary material

**Figure S1.**
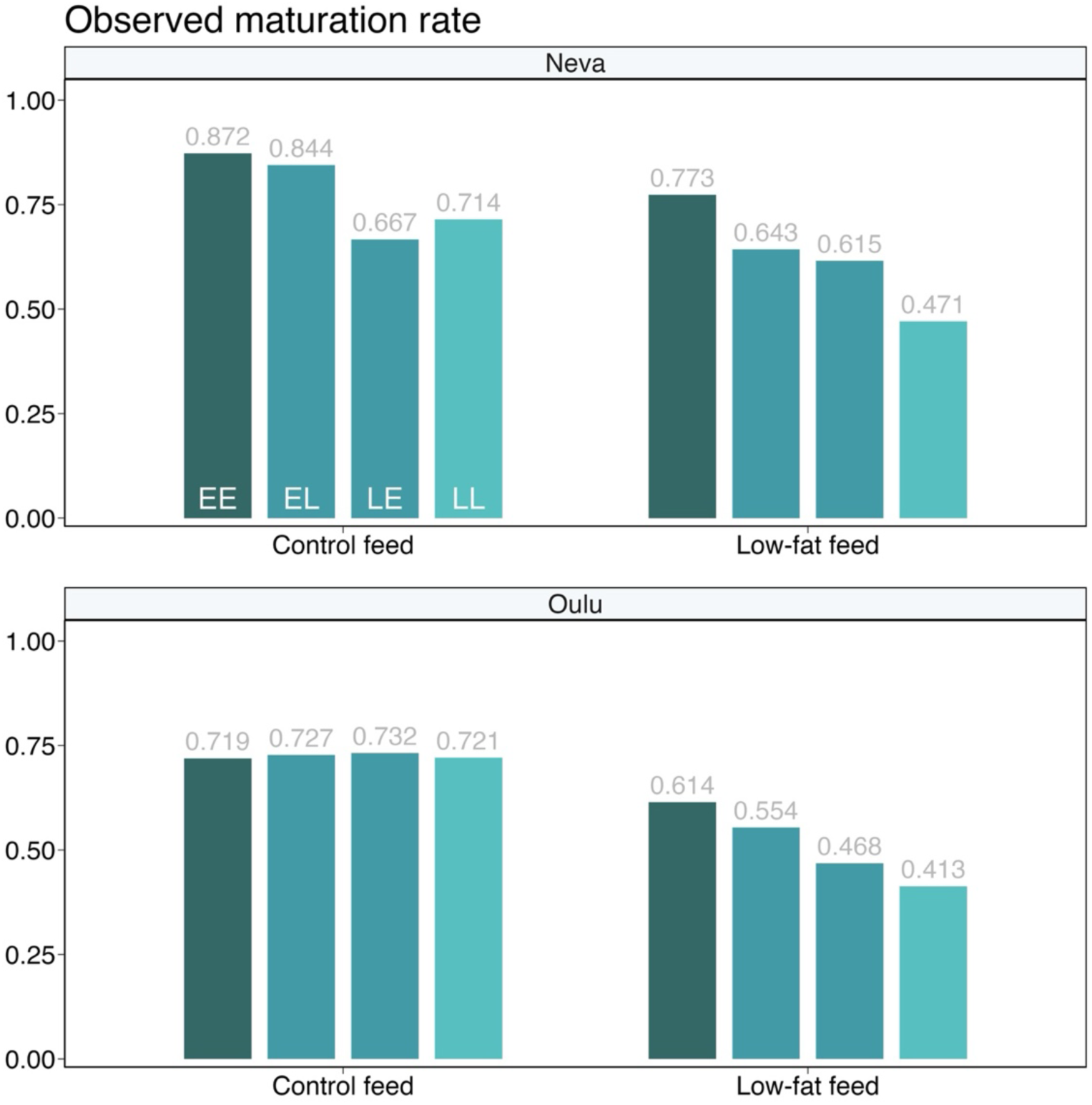
Observed maturation rates of female Atlantic salmon for the Neva and Oulu populations in the two feed treatments (control; low-fat feed). The proportions are shown for each *vgll3* genotype (EE, EL, LE, LL) separately with the differently coloured bars with the exact maturation rate written above each bar. For the heterozygous salmon, the allele inherited from the mother in listed first.

**Table S1.**
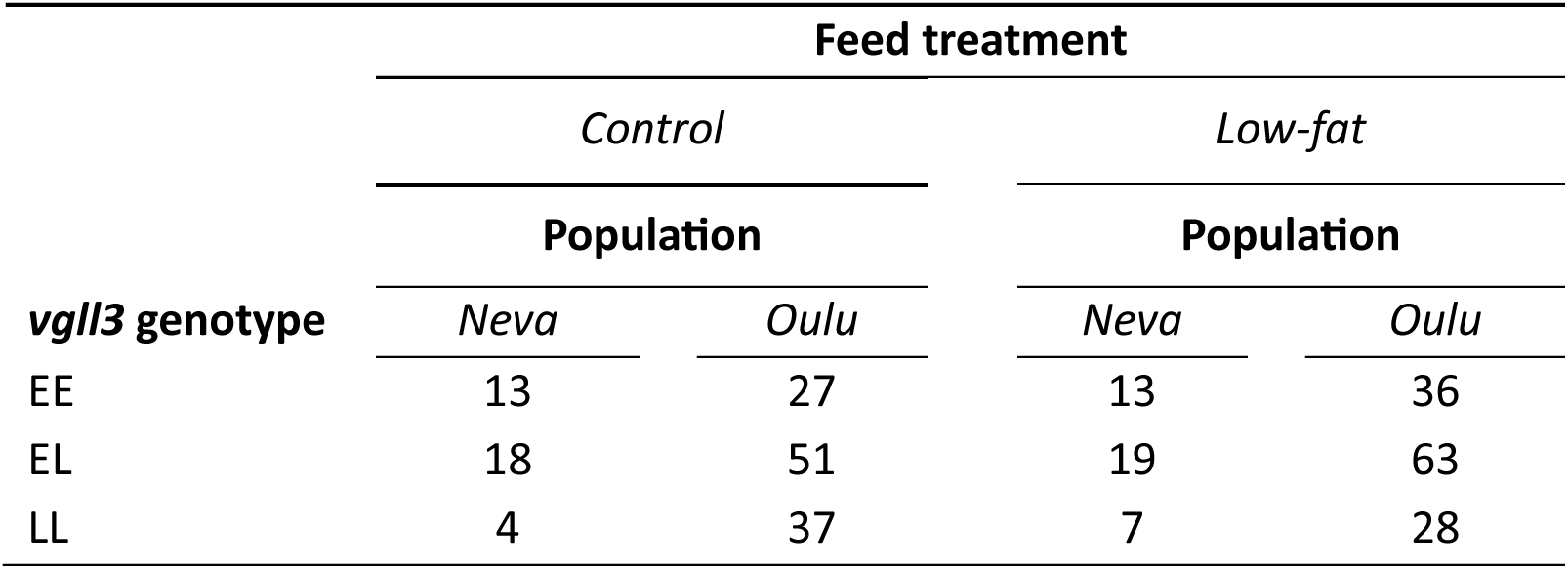
Number of samples with fecundity estimates and their distribution across *vgll3* genotypes, populations, and feed treatments.

**Table S2.** Observed maturation rates for the *vgll3* genotypes (EE, EL, LL) for different treatment categories. The different experimental treatment combinations are listed in the group column. The number within parentheses indicates the number of fish in each treatment combination.

| Group | EE | EL | LL | Overall |
| --- | --- | --- | --- | --- |
| All | 72.9% (*n*=262) | 65.3% (*n*=323) | 59.3% (*n*=145) | 66.8% (*n*=730) |
| Neva | 81.1% (*n*=122) | 72.2% (*n*=115) | 58.1% (*n*=31) | 74.6% (*n*=268) |
| Oulu | 65.7% (*n*=140) | 61.5% (*n*=208) | 59.6% (*n*=114) | 62.3% (*n*=462) |
| Low-fat | 69.0% (*n*=158) | 55.7% (*n*=167) | 42.9% (*n*=63) | 59.0% (*n*=388) |
| Control | 78.8% (*n*=104) | 75.6% (*n*=156) | 72.0% (*n*=82) | 75.7% (*n*=342) |
| Neva, Low-fat | 77.3% (*n*=75) | 63.6% (*n*=55) | 47.1% (*n*=17) | 68.7% (*n*=147) |
| Neva, Control | 87.2% (*n*=47) | 80.0% (*n*=60) | 71.4% (*n*=14) | 81.8% (*n*=121) |
| Oulu, Low-fat | 61.4% (*n*=83) | 51.8% (*n*=112) | 41.3% (*n*=46) | 53.1% (*n*=241) |
| Oulu, Control | 71.9% (*n*=57) | 72.9% (*n*=96) | 72.1% (*n*=68) | 72.4% (*n*=221) |

**Table S3.** Mean length (in cm ± standard error (SE)) for individuals with different *vgll3* genotypes (EE, EL, LL). The different experimental treatment combinations (two populations, two feed treatments) are listed in the group column. Total column indicates maturation rates across all *vgll3* genotypes.

| Group | EE | EL | LL | Total |
| --- | --- | --- | --- | --- |
| Total | 45.7 ± 0.4 | 44.5 ± 0.4 | 43.5 ± 0.5 | 44.8 ± 0.2 |
| Neva | 48.3 ± 0.6 | 48.2 ± 0.6 | 49.4 ± 1.0 | 48.4 ± 0.4 |
| Oulu | 43.4 ± 0.4 | 42.5 ± 0.4 | 42.0 ± 0.5 | 42.6 ± 0.2 |
| Low-fat | 45.1 ± 0.5 | 44.0 ± 0.5 | 42.8 ± 0.9 | 44.3 ± 0.3 |
| Control | 46.5 ± 0.6 | 45.1 ± 0.5 | 44.1 ± 0.6 | 45.3 ± 0.3 |
| Neva, Low-fat | 48.1 ± 0.6 | 47.4 ± 0.8 | 48.4 ± 1.2 | 47.9 ± 0.5 |
| Neva, Control | 48.6 ± 1.0 | 49.0 ± 0.9 | 50.5 ± 1.7 | 49.0 ± 0.6 |
| Oulu, Low-fat | 42.4 ± 0.6 | 42.4 ± 0.5 | 40.8 ± 1.0 | 42.1 ± 0.4 |
| Oulu, Control | 44.8 ± 0.7 | 42.6 ± 0.5 | 42.8 ± 0.6 | 43.2 ± 0.3 |

**Table S4.**
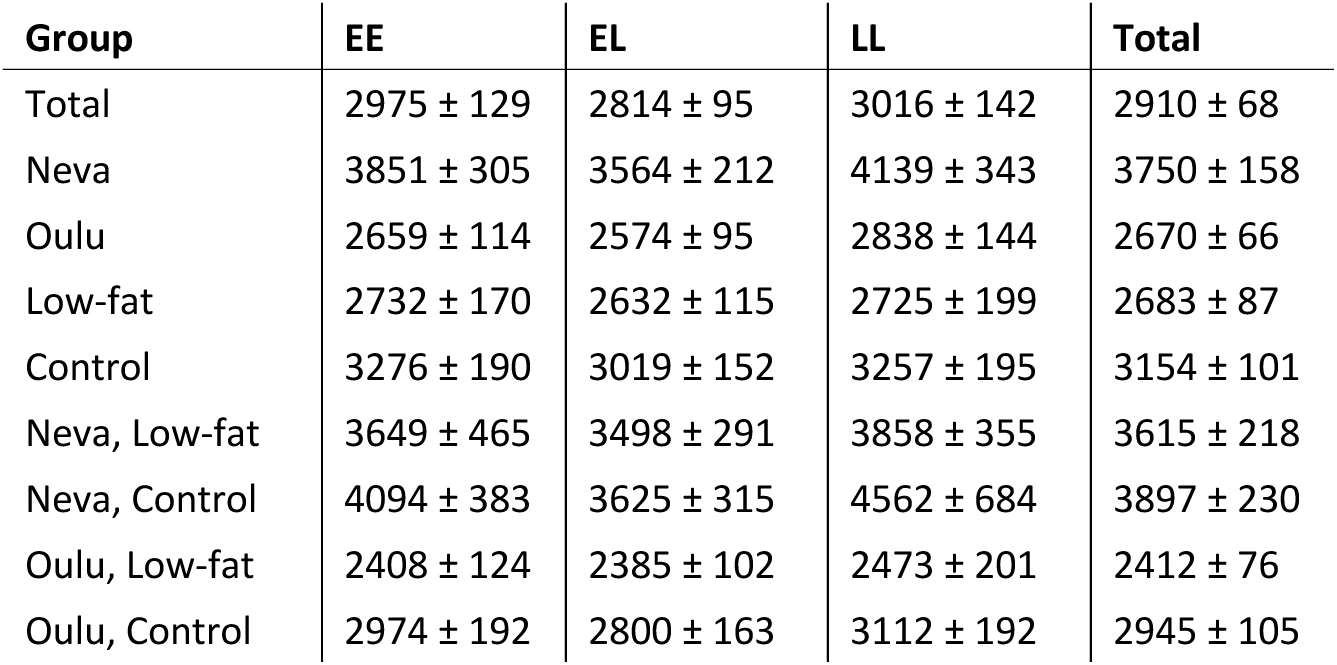
Mean fecundity (total egg number ± SE) for individuals with different *vgll3* genotypes (EE, EL, LL). The different experimental treatment combinations (two populations, two feed treatments) are listed in the group column. Total column indicates maturation rates across all *vgll3* genotypes.

**Figure S2.**
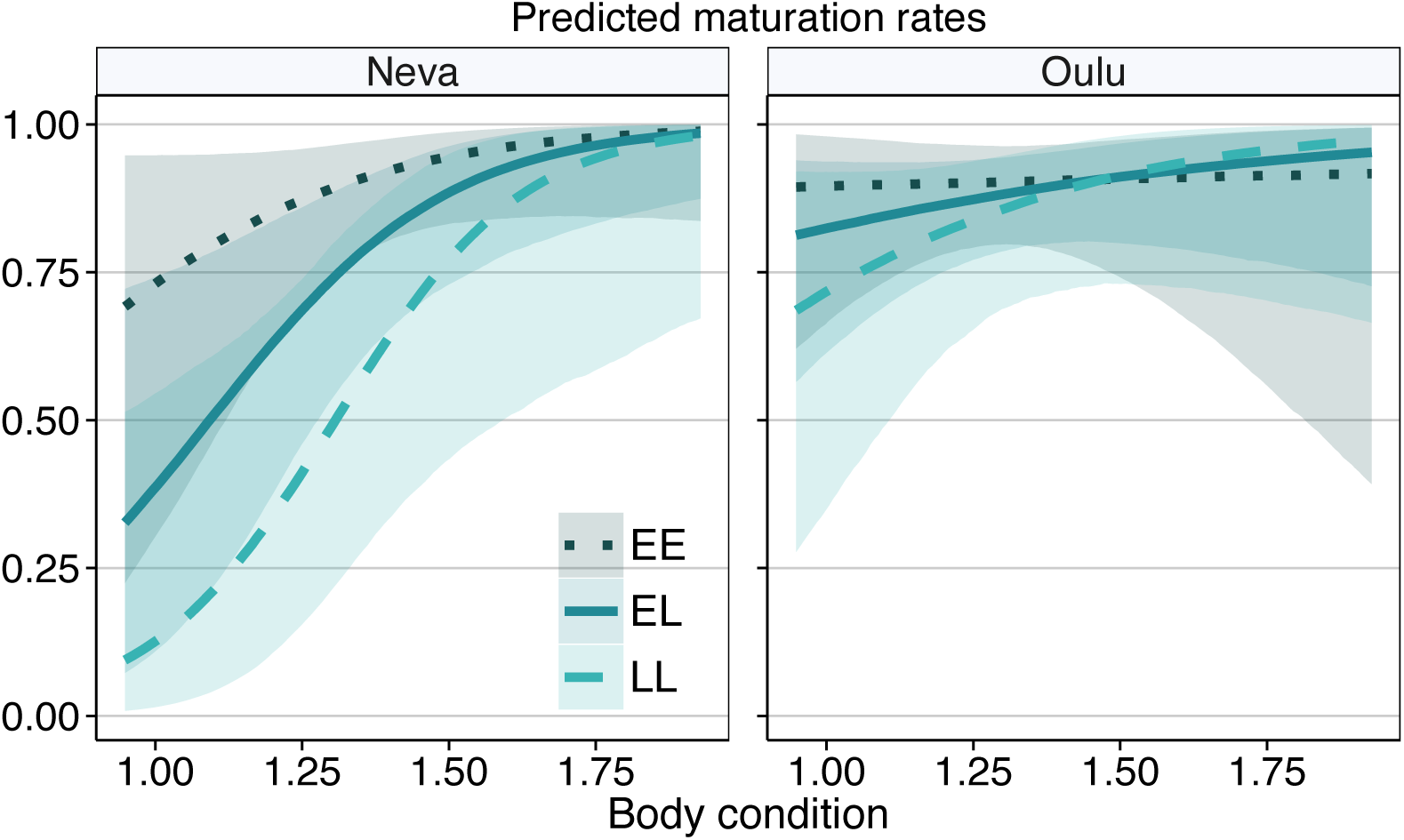
Predicted maturation rate for female Atlantic salmon with different *vgll3* genotypes (dotted dark petrol blue = EE; solid petrol blue = EL; dashed teal = LL) from two different populations (Neva; Oulu) given body condition. Predictions were calculated using the full maturation model with feed treatment set as the low-fat feed and body length as the mean of all the female salmon included in the analysis. The shaded areas along the prediction lines represent the 95% credible intervals.

**Figure S3.**
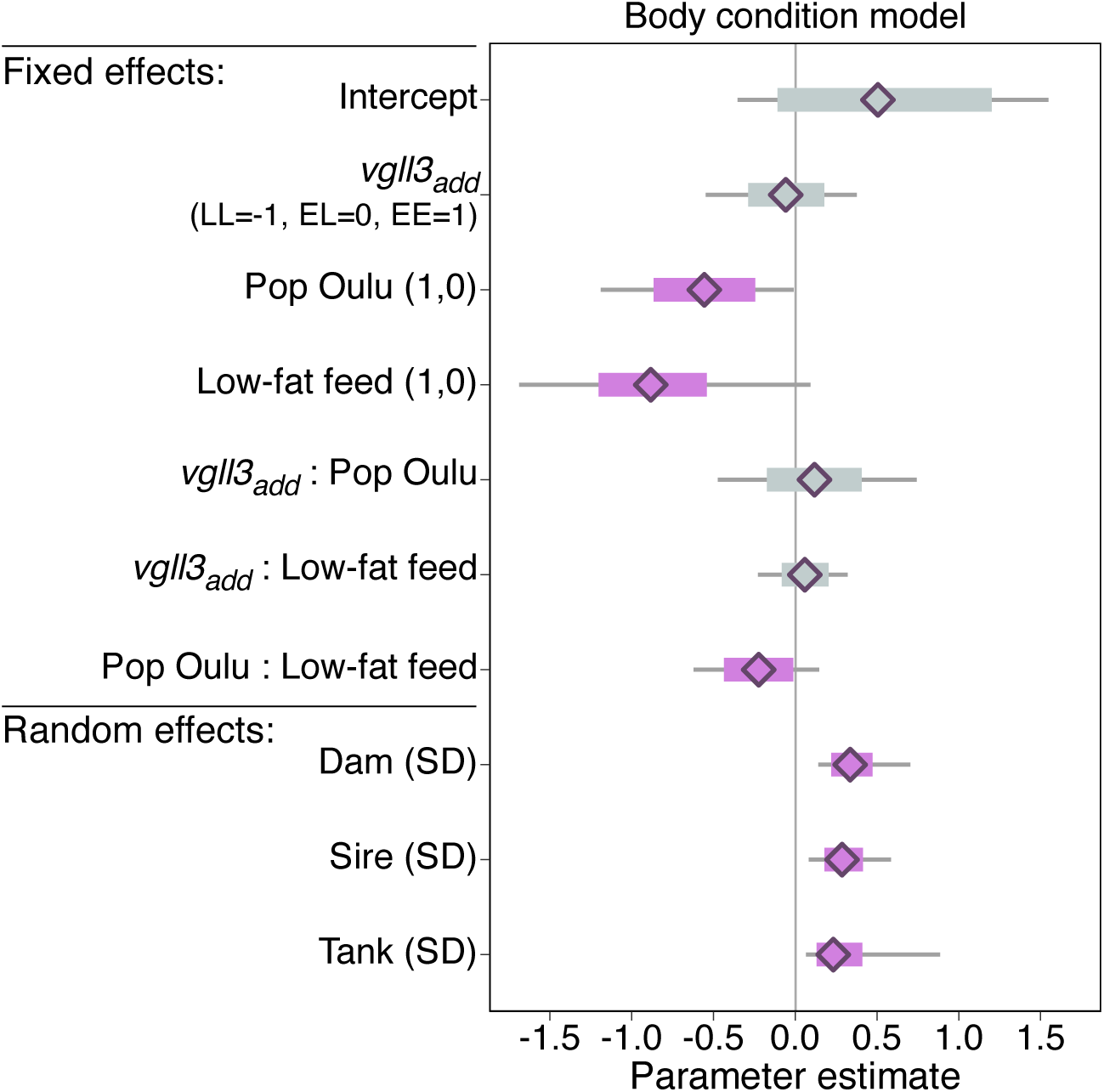
Parameter estimates and 95% credible intervals for the fixed and random effects obtained from the body condition model. The scales of the fixed and random effect model parameters are indicated in the parentheses after the variable name. For categorical variables, the levels are 0 = Neva and 1= Oulu for population, and 0 = control and 1 = low-fat for feed. Continuous variables were centered and standard deviation scaled which leads to model estimates indicating the effect of increasing or decreasing the variable by one standard deviation. Model intercept was set to 0 for all variables. Random effects standard deviations indicate the magnitude of variation between tanks and between parents. The diamonds indicate the mean parameter estimates calculated from the posterior samples, the thick bars indicate the 95% CIs, and the thin bars the 100% CI. The thick bars are coloured purple if the 95% CI does not overlap zero.

**Figure S4.**
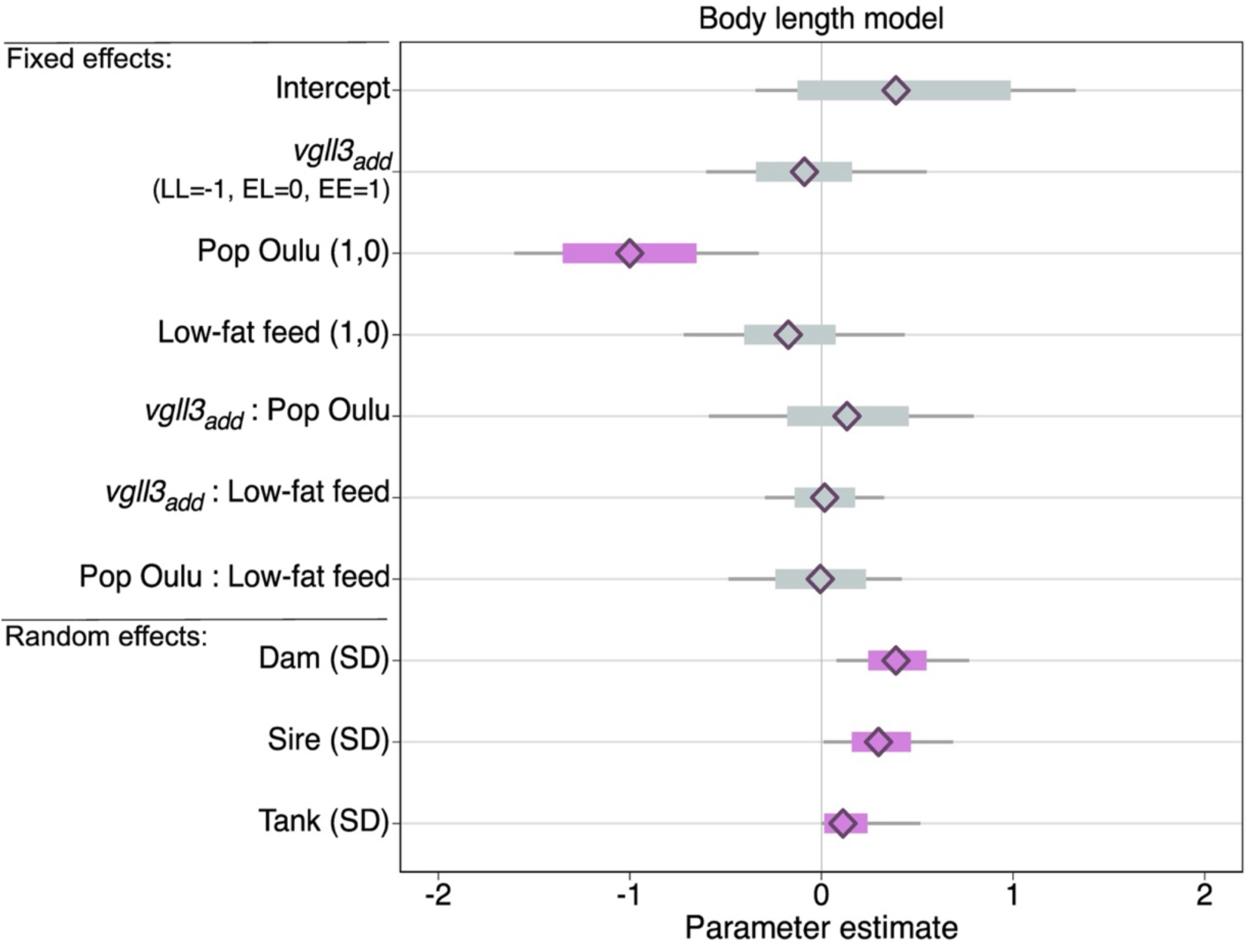
Parameter estimates and 95% credible intervals for the fixed and random effects obtained from the body length model. The scales of the fixed and random effect model parameters are indicated in the parentheses after the variable name. For categorical variables, the levels are 0 = Neva and 1= Oulu for population and 0 = control and 1 = low-fat for feed. Continuous variables were centered and standard deviation scaled which leads to model estimates indicating the effect of increasing or decreasing the variable by one standard deviation. Model intercept was set to 0 for all variables. Random effects standard deviations indicate the magnitude of variation between tanks and between parents. The diamonds indicate the mean parameter estimates calculated from the posterior samples, the thick bars indicate the 95% CIs, and the thin bars the 100% CI. The thick bars are coloured purple if the 95% CI does not overlap zero.

**Figure S5.**
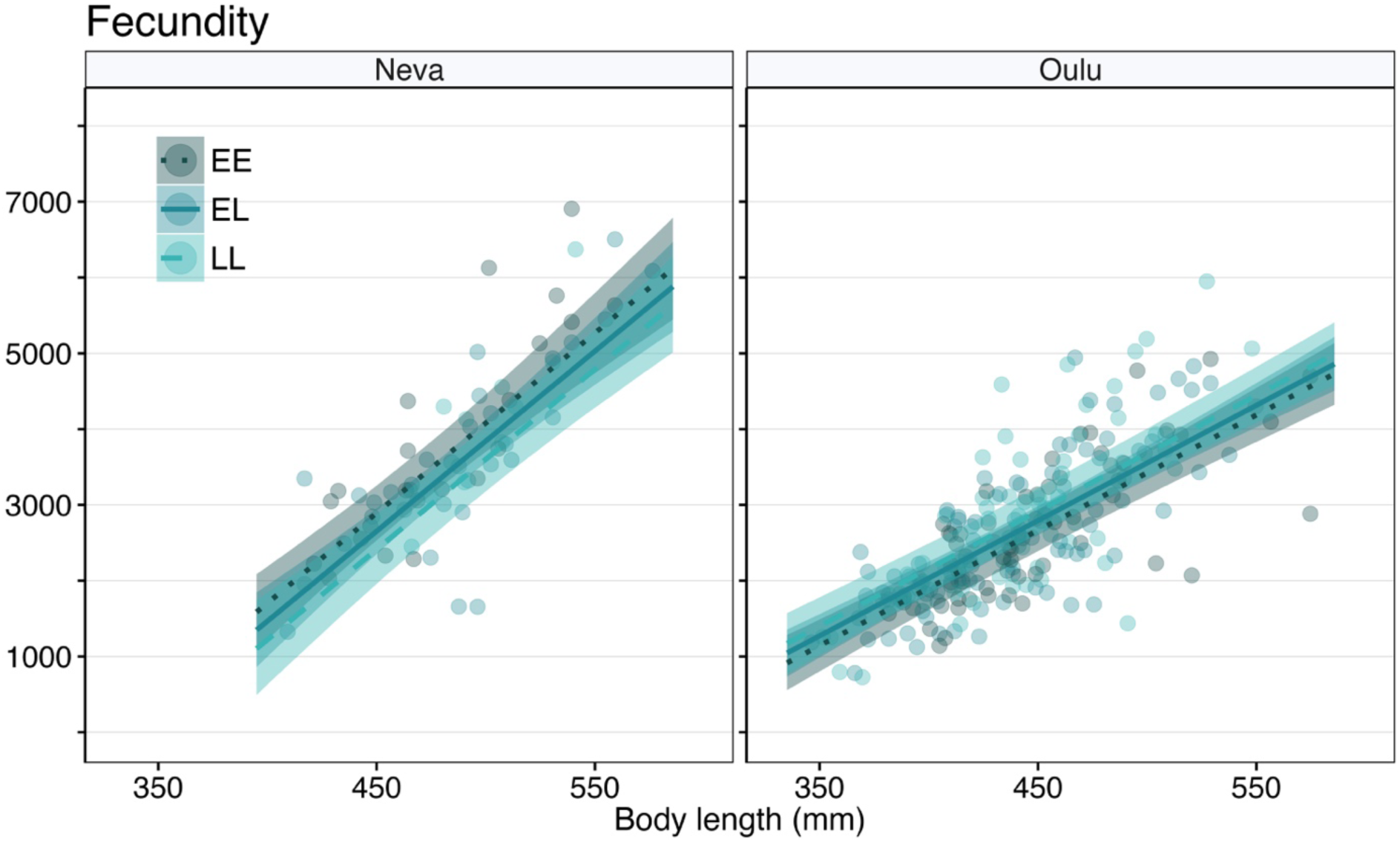
Observed and predicted fecundity (egg number) given body length for two populations (Neva, Oulu) and *vgll3 g*enotypes. Points indicate the observed fecundity for individual females and the trendlines the predicted fecundity based on the full fecundity model. Points and trendlines are coloured based on *vgll3* genotype (dotted dark petrol blue = EE; solid petrol blue = EL; dashed teal = LL).

**Figure S6.**
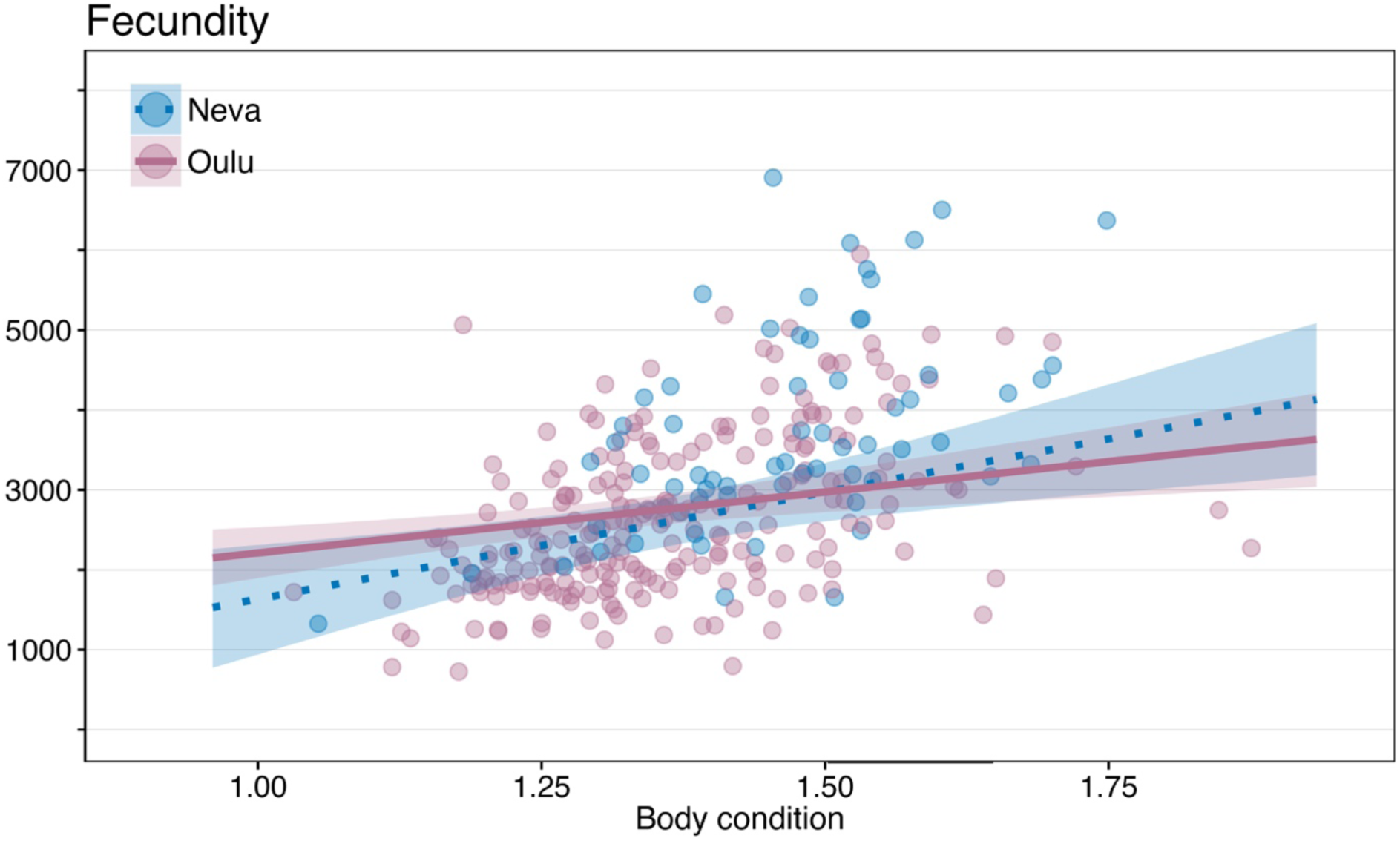
Fecundity (egg number) per mature female plotted against body condition in the spring prior to spawning. The points are the observed fecundity coloured based on the population (blue = Neva, Red = Oulu). The trendlines indicate the predicted fecundity calculated using the fecundity model. For the predictions, *vgll3* genotype was set as EL and feed as low-fat feed. Body length was set as the mean of all the female fish included in the fecundity model. The shaded areas around the prediction lines are the 95% credible intervals.

**Figure S7.**
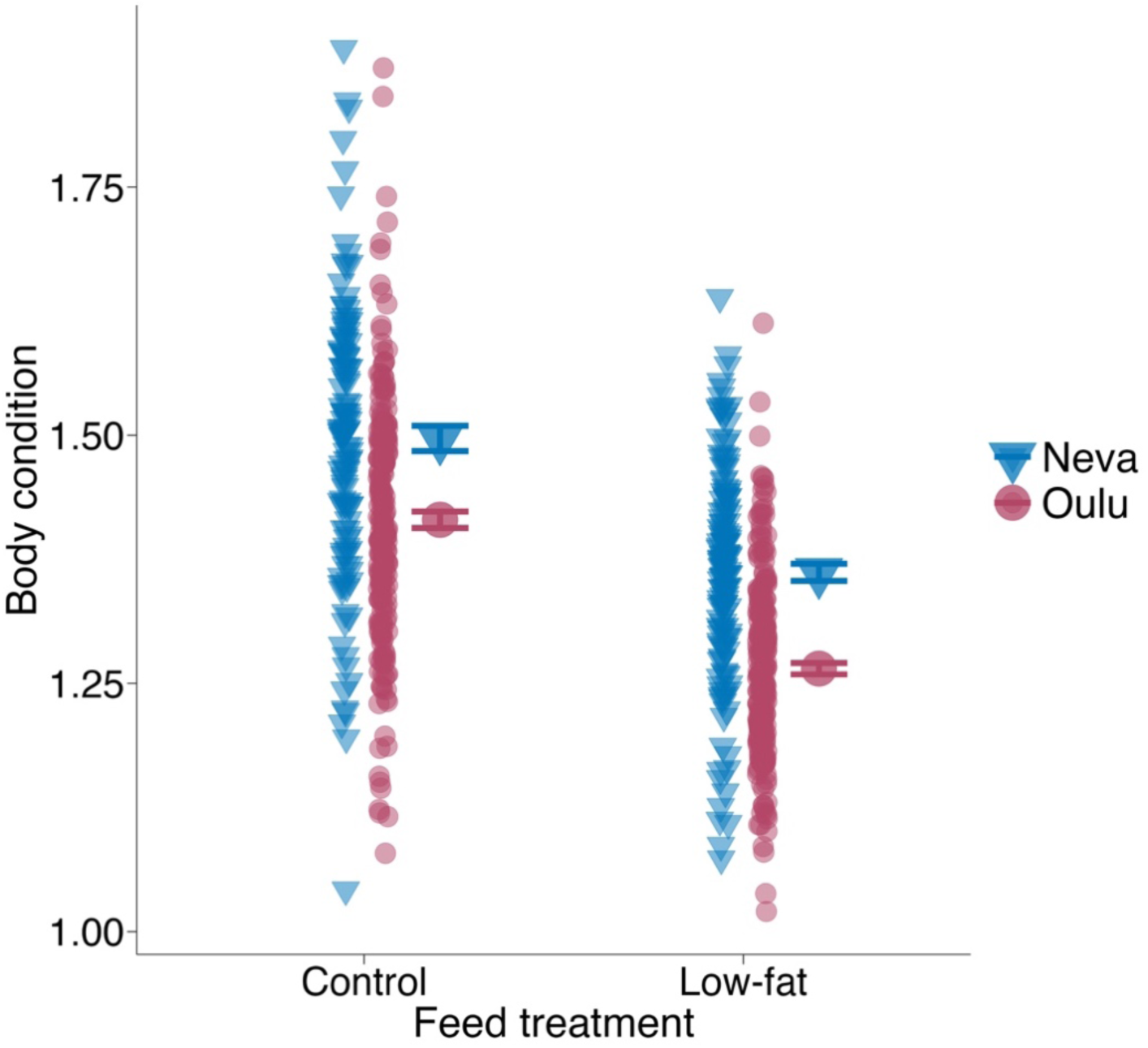
Body condition of female Atlantic salmon in the Control and low-fat feed treatments for the two populations (blue = Neva, red = Oulu). Larger dots with error bars (± SE) indicate the mean body condition and the small dots are the raw data for the females included in the analysis.

**Table S5.** Results of the maturation model. Estimate and Est.Error represent the parameter estimate and its error, respectively. 95% CI (lower) and (upper) are the limits of the 95% credible interval and R-hat, Bulk_ESS, and Tail_ESS (ESS = Estimated sample size) describe model convergence and efficiency.

| **Fixed effects:** | **Estimate** | **Est.Error** | **95% CI (lower)** | **95% CI (upper)** | **R-hat** | **Bulk_ESS** | **Tail_ESS** |
| --- | --- | --- | --- | --- | --- | --- | --- |
| Intercept | 1.58 | 0.49 | 0.65 | 2.57 | 1.00 | 10987 | 7766 |
| *vgll3*<sub>add</sub> (LL=-1, EL=0, EE=1) | 0.88 | 0.41 | 0.07 | 1.69 | 1.00 | 13286 | 8082 |
| Low-fat feed (1,0) | -0.18 | 0.49 | -1.10 | 0.79 | 1.00 | 12786 | 7638 |
| Pop Oulu (1,0) | 0.33 | 0.50 | -0.64 | 1.34 | 1.00 | 15344 | 7854 |
| Condition (SD) | 0.71 | 0.28 | 0.21 | 1.27 | 1.00 | 10860 | 7690 |
| Length (SD) | 1.07 | 0.19 | 0.73 | 1.46 | 1.00 | 9569 | 5416 |
| *vgll3*<sub>add</sub>: Low-fat feed | 0.10 | 0.37 | -0.62 | 0.82 | 1.00 | 15561 | 8619 |
| *vgll3*<sub>add</sub>:Pop Oulu | -0.85 | 0.44 | -1.69 | 0.03 | 1.00 | 12669 | 7686 |
| *vgll3*<sub>add</sub>:Condition | -0.19 | 0.22 | -0.64 | 0.24 | 1.00 | 8858 | 7272 |
| Low-fat feed:Pop Oulu | 0.41 | 0.49 | -0.56 | 1.36 | 1.00 | 14017 | 7983 |
| Pop Oulu:Condition | -0.49 | 0.32 | -1.14 | 0.13 | 1.00 | 9959 | 7506 |
| **Random effects:** |  |  |  |  |  |  |  |
| Animal (SD) | 1.18 | 0.40 | 0.40 | 2.02 | 1.00 | 1491 | 2211 |
| Tank (SD) | 0.45 | 0.23 | 0.07 | 0.96 | 1.00 | 3054 | 3707 |

**Table S6.** Results of the body condition model. Estimate and Est.Error represent the parameter estimate and its error, respectively. 95% CI (lower) and (upper) are the limits of the 95% credible interval and R-hat, Bulk_ESS, and Tail_ESS (ESS = Estimated sample size) describe model convergence and efficiency.

| **Fixed effects:** | **Estimate** | **Est.Error** | **95% CI (lower)** | **95% CI (upper)** | **R-hat** | **Bulk_ESS** | **Tail_ESS** |
| --- | --- | --- | --- | --- | --- | --- | --- |
| Intercept | 0.95 | 0.15 | 0.65 | 1.25 | 1.00 | 6806 | 6644 |
| *vgll3*<sub>add</sub> (LL=-1, EL=0, EE=1) | -0.06 | 0.12 | -0.29 | 0.18 | 1.00 | 7785 | 7446 |
| Pop Oulu (1,0) | -0.56 | 0.16 | -0.87 | -0.25 | 1.00 | 7744 | 6973 |
| Low-fat feed (1,0) | -0.88 | 0.17 | -1.2 | -0.54 | 1.00 | 5936 | 6265 |
| *vgll3*<sub>add</sub>:Pop Oulu | 0.12 | 0.15 | -0.17 | 0.41 | 1.00 | 7772 | 7500 |
| *vgll3*<sub>add</sub>: Low-fat feed | 0.06 | 0.07 | -0.08 | 0.2 | 1.00 | 19143 | 7533 |
| Low-fat feed:Pop Oulu | -0.22 | 0.11 | -0.44 | -0.01 | 1.00 | 17883 | 7934 |
| **Random effects:** |  |  |  |  |  |  |  |
| Dam (SD) | 0.34 | 0.06 | 0.22 | 0.47 | 1.00 | 4093 | 6407 |
| Sire (SD) | 0.29 | 0.06 | 0.18 | 0.41 | 1.00 | 3427 | 6203 |
| Tank (SD) | 0.23 | 0.07 | 0.13 | 0.41 | 1.00 | 4291 | 6385 |

**Table S7.** Results of the body length model. Estimate and Est.Error represent the parameter estimate and its error, respectively. 95% CI (lower) and (upper) are the limits of the 95% credible interval and R-hat, Bulk_ESS, and Tail_ESS (ESS = Estimated sample size) describe model convergence and efficiency.

| **Fixed effects:** | **Estimate** | **Est.Error** | **95% CI**<br>**(lower)** | **95% CI**<br>**(upper)** | **R-hat** | **Bulk_ESS** | **Tail_ESS** |
| --- | --- | --- | --- | --- | --- | --- | --- |
| Intercept | 0.76 | 0.14 | 0.48 | 1.04 | 1.00 | 9078 | 6004 |
| *vgll3*<sub>add</sub> (LL=-1, EL=0, EE=1) | -0.09 | 0.13 | -0.34 | 0.16 | 1.00 | 8264 | 7236 |
| Pop Oulu (1,0) | -1.00 | 0.18 | -1.35 | -0.65 | 1.00 | 7775 | 7213 |
| Low-fat feed (1,0) | -0.17 | 0.12 | -0.40 | 0.07 | 1.00 | 9763 | 7264 |
| *vgll3*<sub>add</sub>:Pop Oulu | 0.13 | 0.16 | -0.18 | 0.46 | 1.00 | 7134 | 7014 |
| *vgll3*<sub>add</sub>: Low-fat feed | 0.02 | 0.08 | -0.14 | 0.18 | 1.00 | 21167 | 7842 |
| Low-fat feed:Pop Oulu | -0.01 | 0.12 | -0.24 | 0.23 | 1.00 | 13653 | 8090 |
| **Random effects:** |  |  |  |  |  |  |  |
| Dam (SD) | 0.39 | 0.08 | 0.24 | 0.55 | 1.00 | 2859 | 4958 |
| Sire (SD) | 0.30 | 0.08 | 0.16 | 0.47 | 1.00 | 2051 | 3684 |
| Tank (SD) | 0.11 | 0.06 | 0.01 | 0.24 | 1.00 | 2837 | 3372 |

**Table S8.** Results of the fecundity model. Estimate and Est.Error represent the parameter estimate and its error, respectively. 95% CI (lower) and (upper) are the limits of the 95% credible interval and R-hat, Bulk_ESS, and Tail_ESS (ESS = Estimated sample size) describe model convergence and efficiency.

| **Fixed effects:** | **Estimate** | **Est.Error** | **95% CI**<br>**(lower)** | **95% CI**<br>**(upper)** | **R-hat** | **Bulk_ESS** | **Tail_ESS** |
| --- | --- | --- | --- | --- | --- | --- | --- |
| Intercept | -0.35 | 0.16 | -0.65 | -0.04 | 1.00 | 5973 | 7330 |
| *vgll3*<sub>add</sub> (LL=-1, EL=0, EE=1) | 0.18 | 0.16 | -0.12 | 0.49 | 1.00 | 3943 | 5294 |
| Pop Oulu (1,0) | 0.31 | 0.17 | -0.03 | 0.66 | 1.00 | 5932 | 6983 |
| Low-fat feed (1,0) | 0.11 | 0.15 | -0.19 | 0.41 | 1.00 | 5865 | 7065 |
| Condition (SD) | 0.32 | 0.1 | 0.12 | 0.52 | 1.00 | 5013 | 6741 |
| Length (SD) | 1.13 | 0.12 | 0.9 | 1.35 | 1.00 | 5854 | 6668 |
| *vgll3*<sub>add</sub>:Pop Oulu | -0.32 | 0.14 | -0.61 | -0.04 | 1.00 | 6296 | 7295 |
| Low-fat feed:Pop Oulu | -0.19 | 0.17 | -0.53 | 0.15 | 1.00 | 6008 | 7090 |
| *vgll3*<sub>add</sub>:Low-fat feed | 0.03 | 0.11 | -0.2 | 0.26 | 1.00 | 7294 | 7723 |
| *vgll3*<sub>add</sub>:Condition | 0.01 | 0.06 | -0.12 | 0.13 | 1.00 | 6159 | 7499 |
| Pop Oulu:Condition | -0.14 | 0.12 | -0.36 | 0.09 | 1.00 | 5018 | 6865 |
| Pop Oulu:Length | -0.41 | 0.13 | -0.66 | -0.16 | 1.00 | 5753 | 6974 |
| **Random effects:** |  |  |  |  |  |  |  |
| Dam (SD) | 0.14 | 0.08 | 0.01 | 0.30 | 1.00 | 1429 | 3529 |
| Sire (SD) | 0.18 | 0.08 | 0.03 | 0.33 | 1.00 | 1253 | 1480 |
| Tank (SD) | 0.05 | 0.04 | 0 | 0.14 | 1.00 | 5526 | 4902 |

## References

Åsheim, E. R., Debes, P. V., House, A., Liljeström, P., Niemelä, P. T., Siren, J. P., Erkinaro, J., & Primmer, C. R. (2023). Atlantic salmon (Salmo salar) age at maturity is strongly affected by temperature, population and age-at-maturity genotype. Conservation Physiology, 11(1), coac086. 10.1093/conphys/coac086

Ayllon, F., Kjærner-Semb, E., Furmanek, T., Wennevik, V., Solberg, M. F., Dahle, G., Taranger, G. L., Glover, K. A., Almén, M. S., Rubin, C. J., Edvardsen, R. B., & Wargelius, A. (2015). The vgll3 Locus Controls Age at Maturity in Wild and Domesticated Atlantic Salmon (Salmo salar L.) Males. PLoS Genetics, 11(11), 1–15. 10.1371/journal.pgen.1005628

Ayllon, F., Solberg, M. F., Glover, K. A., Mohammadi, F., Kjærner-Semb, E., Fjelldal, P. G., Andersson, E., Hansen, T., Edvardsen, R. B., & Wargelius, A. (2019). The influence of vgll3 genotypes on sea age at maturity is altered in farmed mowi strain Atlantic salmon. BMC Genetics, 20(1), 1–8. 10.1186/s12863-019-0745-9

Bangura, P. B., Tiira, K., Aykanat, T., Niemelä, P. T., Erkinaro, J., Liljeström, P., Toikkanen, A., & Primmer, C. R. (2024). Sex-specific associations of the maturation locus vgll3 with exploratory behavior and boldness in Atlantic salmon juveniles. Ecology and Evolution, 14(6), e11449. 10.1002/ece3.11449

Bangura, P. B., Tiira, K., Niemelä, P. T., Erkinaro, J., Liljeström, P., Toikkanen, A., & Primmer, C. R. (2022). Linking vgll3 genotype and aggressive behaviour in juvenile Atlantic salmon (Salmo salar). Journal of Fish Biology, 100(5), 1264–1271. 10.1111/jfb.15040

Barneche, D. R., Robertson, D. R., White, C. R., & Marshall, D. J. (2018). Fish reproductive-energy output increases disproportionately with body size. Science, 360(6389), 642– 645. 10.1126/science.aao6868

Barson, N. J., Aykanat, T., Hindar, K., Baranski, M., Bolstad, G. H., Fiske, P., Jacq, C., Jensen, A. J., Johnston, S. E., Karlsson, S., Kent, M., Moen, T., Niemelä, E., Nome, T., Næsje, T. F., Orell, P., Romakkaniemi, A., Sægrov, H., Urdal, K., … Primmer, C. R. (2015). Sex-dependent dominance at a single locus maintains variation in age at maturity in salmon. Nature, 528(7582), 405–408. 10.1038/nature16062

Bernardo, J. (1993). Determinants of maturation in animals. Trends in Ecology & Evolution, 8(5), 166–173. 10.1016/0169-5347(93)90142-C

Besnier, F., Skaala, Ø., Wennevik, V., Ayllon, F., Utne, K. R., Fjeldheim, P. T., Andersen-Fjeldheim, K., Knutar, S., & Glover, K. A. (2024). Overruled by nature: A plastic response to environmental change disconnects a gene and its trait. Molecular Ecology, 33(2), e16933. 10.1111/mec.16933

Braendle, C., Heyland, A., & Flatt, T. (2011). Integrating mechanistic and evolutionary analysis of life history variation. In T. Flatt & A. Heyland (Eds.), Mechanisms of Life History Evolution (pp. 3–10). Oxford University Press.

Bürkner, P.-C. (2017). brms: An R Package for Bayesian Multilevel Models Using Stan. Journal of Statistical Software, 80(1). 10.18637/jss.v080.i01

Bürkner, P.-C. (2018). Advanced Bayesian Multilevel Modeling with the R Package brms. The R Journal, 10(1), 395. 10.32614/RJ-2018-017

Bürkner, P.-C. (2021). Bayesian Item Response Modeling in *R* with **brms** and *Stan*. Journal of Statistical Software, 100(5). 10.18637/jss.v100.i05

Carlson, S. M., & Seamons, T. R. (2008). A review of quantitative genetic components of fitness in salmonids: Implications for adaptation to future change. Evolutionary Applications, 1(2), 222–238. 10.1111/j.1752-4571.2008.00025.x

Chaput, G., Allard, J., Caron, F., Dempson, J. B., Mullins, C. C., & O’Connell, M. F. (1998). River-specific target spawning requirements for Atlantic salmon (Salmo salar) based on a generalized smolt production model. Canadian Journal of Fisheries and Aquatic Sciences, 55(1), 246–261. 10.1139/f97-252

Cheng, S., Jacobs, C. G. C., Mogollón Pérez, E. A., Chen, D., van de Sanden, J. T., Bretscher, K. M., Verweij, F., Bosman, J. S., Hackmann, A., Merks, R. M. H., van den Heuvel, J., & van der Zee, M. (2024). A life-history allele of large effect shortens developmental time in a wild insect population. Nature Ecology & Evolution, 8(1), 70–82. 10.1038/s41559-023-02246-y

Czorlich, Y., Aykanat, T., Erkinaro, J., Orell, P., & Primmer, C. R. (2018). Rapid sex-specific evolution of age at maturity is shaped by genetic architecture in Atlantic salmon. Nature Ecology and Evolution, 2(11), 1800–1807. 10.1038/s41559-018-0681-5

de Eyto, E., White, J., Boylan, P., Clarke, B., Cotter, D., Doherty, D., Gargan, P., Kennedy, R., McGinnity, P., O’Maoiléidigh, N., & O’Higgins, K. (2015). The fecundity of wild Irish Atlantic salmon Salmo salar L. and its application for stock assessment purposes. Fisheries Research, 164, 159–169. 10.1016/j.fishres.2014.11.017

Debes, P. V., Piavchenko, N., Erkinaro, J., & Primmer, C. R. (2020). Genetic growth potential, rather than phenotypic size, predicts migration phenotype in Atlantic salmon. Proceedings of the Royal Society B: Biological Sciences, 287(1931). 10.1098/rspb.2020.0867rspb20200867

Debes, P. V., Piavchenko, N., Ruokolainen, A., Ovaskainen, O., Moustakas-Verho, J. E., Parre, N., Aykanat, T., Erkinaro, J., & Primmer, C. R. (2021). Polygenic and major-locus contributions to sexual maturation timing in Atlantic salmon. Molecular Ecology, 00, 1–15. 10.1111/mec.16062

Erkinaro, J., Czorlich, Y., Orell, P., Kuusela, J., Falkegård, M., Länsman, M., Pulkkinen, H., Primmer, C. R., & Niemelä, E. (2019). Life history variation across four decades in a diverse population complex of atlantic salmon in a large subarctic river. Canadian Journal of Fisheries and Aquatic Sciences, 76(1), 42–55. 10.1139/cjfas-2017-0343

Evans, M. L., Hard, J. J., Black, A. N., Sard, N. M., & O’Malley, K. G. (2019). A quantitative genetic analysis of life-history traits and lifetime reproductive success in reintroduced Chinook salmon. Conservation Genetics, 20(4), 781–799. 10.1007/s10592-019-01174-4

Falconer, D. S., & Mackay, T. F. C. (1996). Introduction to quantitative genetics (4th ed.). Longman.

Fleming, I. A. (1996). Reproductive strategies of Atlantic salmon: Ecology and evolution. Reviews in Fish Biology and Fisheries, 6(4), 379–416. 10.1007/BF00164323

Fleming, I. A., & Einum, S. (2011). Reproductive Ecology: A Tale of Two Sexes. In Ø. Aas, S. Einum, A. Klemetsen, & J. Skurdal (Eds.), Atlantic Salmon Ecology. Blackwell Publishing Ltd.

Fleming, I. A., Jonsson, B., Gross, M. R., & Lamberg, A. (1996). An Experimental Study of the Reproductive Behaviour and Success of Farmed and Wild Atlantic Salmon (Salmo salar). Journal of Applied Ecology, 33(4), 893–905. 10.2307/2404960

Forseth, T., Fiske, P., Barlaup, B., Gjøsæter, H., Hindar, K., & Diserud, O. H. (2013). Reference point based management of Norwegian Atlantic salmon populations. Environmental Conservation, 40(4), 356–366. 10.1017/S0376892913000416

Ginther, S. C., Cameron, H., White, C. R., & Marshall, D. J. (2024). Metabolic loads and the costs of metazoan reproduction. Science, 384(6697), 763–767. 10.1126/science.adk6772

Hanson, N., Ounsley, J., Burton, T., Auer, S., Hunt, J. H., Shaw, B., Henderson, J., & Middlemas, S. J. (2020). Hierarchical analysis of wild Atlantic salmon (Salmo salar) fecundity in relation to body size and developmental traits. Journal of Fish Biology, 96(2), 316–326. 10.1111/jfb.14181

Heincke, F. (1908). Bericht über die Untersuchungen der Biologischen Anstalt auf Helgoland zur Naturgeschichte der Nutzfische (IV/V. Bericht Über Die Beteiligung Deutschlands an Der Internationalen Meeresforschung in Den Jahren 1905/6–1906/7, pp. 67–156). Deutsche Wissenschaftliche Kommission für Meeresforschung.

Hess, J. E., Zendt, J. S., Matala, A. R., & Narum, S. R. (2016). Genetic basis of adult migration timing in anadromous steelhead discovered through multivariate association testing. Proceedings of the Royal Society B: Biological Sciences, 283(1830), 20153064. 10.1098/rspb.2015.3064

Hixon, M. A., Johnson, D. W., & Sogard, S. M. (2014). BOFFFFs: On the importance of conserving old-growth age structure in fishery populations. ICES Journal of Marine Science, 71(8), 2171–2185. 10.1093/icesjms/fst200

House, A. H., Debes, P. V., Kurko, J., Erkinaro, J., & Primmer, C. R. (2023). Genotype-specific variation in seasonal body condition at a large-effect maturation locus. Proceedings of the Royal Society B: Biological Sciences, 290(1999), 20230432. 10.1098/rspb.2023.0432

Hutchings, J. A. (2021). A Primer of Life Histories: Ecology, Evolution, and Application (1st ed.). Oxford University Press. 10.1093/oso/9780198839873.001.0001

Hutchings, J. A., & Jones, M. E. B. (1998). Life history variation and growth rate thresholds for maturity in Atlantic salmon, Salmo salar. Canadian Journal of Fisheries and Aquatic Sciences, 55(SUPPL.1), 22–47. 10.1139/cjfas-55-s1-22

Jacobson, P., Whitlock, R., Huss, M., Leonardsson, K., Östergren, J., & Gårdmark, A. (2021). Growth variation of Atlantic salmon Salmo salar at sea affects their population-specific reproductive potential. Marine Ecology Progress Series, 671, 165–174. 10.3354/meps13734

Jonsson, B., Finstad, A. G., & Jonsson, N. (2012). Winter temperature and food quality affect age at maturity: An experimental test with Atlantic salmon (Salmo salar). Canadian Journal of Fisheries and Aquatic Sciences, 69(11), 1817–1826. 10.1139/f2012-108

Jonsson, B., & Jonsson, N. (2011). Ecology of Atlantic Salmon and Brown Trout—Habitat as a Template for Life Histories. Springer. 10.1007/978-94-007-1189-1

Jonsson, B., Jonsson, N., & Finstad, A. G. (2013). Effects of temperature and food quality on age and size at maturity in ectotherms: An experimental test with Atlantic salmon. Journal of Animal Ecology, 82(1), 201–210. 10.1111/j.1365-2656.2012.02022.x

Kadri, S., Metcalfe, N. B., Huntingford, F. A., Thorpe, J. E., & Mitchell, D. F. (1997). Early morphological predictors of maturity in one-sea-winter Atlantic salmon. Aquaculture International, 5(1), 41–50. 10.1007/BF02764786

Klemetsen, A., Amundsen, P. A., Dempson, J. B., Jonsson, B., Jonsson, N., O’Connell, M. F., & Mortensen, E. (2003). Atlantic salmon Salmo salar L., brown trout Salmo trutta L. and Arctic charr Salvelinus alpinus (L.): A review of aspects of their life histories. Ecology of Freshwater Fish, 12(1), 1–59. 10.1034/j.1600-0633.2003.00010.x

Kuparinen, A., & Hutchings, J. A. (2017). Genetic architecture of age at maturity can generate divergent and disruptive harvest-induced evolution. Philosophical Transactions of the Royal Society B: Biological Sciences, 372(1712), 20160035. 10.1098/rstb.2016.0035

Küpper, C., Stocks, M., Risse, J. E., dos Remedios, N., Farrell, L. L., McRae, S. B., Morgan, T. C., Karlionova, N., Pinchuk, P., Verkuil, Y. I., Kitaysky, A. S., Wingfield, J. C., Piersma, T., Zeng, K., Slate, J., Blaxter, M., Lank, D. B., & Burke, T. (2016). A supergene determines highly divergent male reproductive morphs in the ruff. Nature Genetics, 48(1), 79–83. 10.1038/ng.3443

Lamichhaney, S., Fan, G., Widemo, F., Gunnarsson, U., Thalmann, D. S., Hoeppner, M. P., Kerje, S., Gustafson, U., Shi, C., Zhang, H., Chen, W., Liang, X., Huang, L., Wang, J., Liang, E., Wu, Q., Lee, S. M.-Y., Xu, X., Höglund, J., … Andersson, L. (2016). Structural genomic changes underlie alternative reproductive strategies in the ruff (Philomachus pugnax). Nature Genetics, 48(1), 84–88. 10.1038/ng.3430

Lynch, M., & Walsh, B. (1998). Genetics and Analysis of Quantitative Traits. Sinauer Associates.

Maamela, K. S., Åsheim, E. R., Debes, P. V., House, A. H., Erkinaro, J., Liljeström, P., Primmer, C. R., & Mobley, K. B. (2023). The effect of temperature and dietary energy content on female maturation and egg nutritional content in Atlantic salmon. Journal of Fish Biology, 102(5), 1096–1108. 10.1111/jfb.15318

Marshall, D. J., Bode, M., Mangel, M., Arlinghaus, R., & Dick, E. J. (2021). Reproductive hyperallometry and managing the world’s fisheries. Proceedings of the National Academy of Sciences, 118(34), e2100695118. 10.1073/pnas.2100695118

Miettinen, A., Romakkaniemi, A., Dannewitz, J., Pakarinen, T., Palm, S., Persson, L., Östergren, J., Primmer, C. R., & Pritchard, V. L. (2024). Temporal allele frequency changes in large-effect loci reveal potential fishing impacts on salmon life-history diversity. Evolutionary Applications, 17(4), e13690. 10.1111/eva.13690

Mobley, K. B., Aykanat, T., Czorlich, Y., House, A., Kurko, J., Miettinen, A., Moustakas-Verho, J. E., Salgado, A., Sinclair-Waters, M., Verta, J.-P., & Primmer, C. R. (2021). Maturation in Atlantic salmon (Salmo salar, Salmonidae): A review of ecological, genetic, and molecular processes. Rev Fish Biol Fisheries, 31, 523–571. 10.1007/s11160-021-09656-w

Mobley, K. B., Barton, H. J., Ellmén, M., Ruokolainen, A., Guttorm, O., Pieski, H., Orell, P., Erkinaro, J., & Primmer, C. R. (2024). Sex-specific overdominance at the maturation vgll3 gene for reproductive fitness in wild Atlantic salmon. Molecular Ecology, 33(14), e17435. 10.1111/mec.17435

Mobley, K. B., Granroth-Wilding, H., Ellmén, M., Orell, P., Erkinaro, J., & Primmer, C. R. (2020). Time spent in distinct life history stages has sex-specific effects on reproductive fitness in wild Atlantic salmon. Molecular Ecology, 29(6), 1173–1184. 10.1111/mec.15390

Moffett, I. J. J., Allen, M., Flanagan, C., Crozier, W. W., & Kennedy, G. J. A. (2006). Fecundity, egg size and early hatchery survival for wild Atlantic salmon, from the River Bush. Fisheries Management and Ecology, 13(2), 73–79. 10.1111/j.1365-2400.2006.00478.x

Mousseau, T. A., & Roff, D. A. (1987). Natural selection and the heritability of fitness components. Heredity, 59(2), 181–197. 10.1038/hdy.1987.113

Niemelä, P. T., Klemme, I., Karvonen, A., Hyvärinen, P., Debes, P. V., Erkinaro, J., Sinclair-Waters, M., Pritchard, V. L., Härkönen, L. S., & Primmer, C. R. (2022). Life-history genotype explains variation in migration activity in Atlantic salmon (Salmo salar). Proceedings of the Royal Society B. 10.1098/rspb.2022.0851

O’Connell, M. F., & Dempson, J. B. (1995). Target spawning requirements for Atlantic salmon, Salmo salar L., in Newfoundland rivers. Fisheries Management and Ecology, 2(3), 161–170. 10.1111/j.1365-2400.1995.tb00109.x

Oomen, R. A., & Hutchings, J. A. (2022). Genomic reaction norms inform predictions of plastic and adaptive responses to climate change. Journal of Animal Ecology, 91(6), 1073–1087. 10.1111/1365-2656.13707

Oomen, R. A., Kuparinen, A., & Hutchings, J. A. (2020). Consequences of Single-Locus and Tightly Linked Genomic Architectures for Evolutionary Responses to Environmental Change. Journal of Heredity, 111(4), 319–332. 10.1093/jhered/esaa020

Persson, L., Raunsgard, A., Thorstad, E. B., Østborg, G., Urdal, K., Sægrov, H., Ugedal, O., Hindar, K., Karlsson, S., Fiske, P., & Bolstad, G. H. (2023). Iteroparity and its contribution to life-history variation in Atlantic salmon. Canadian Journal of Fisheries and Aquatic Sciences, 80(3), 577–592. 10.1139/cjfas-2022-0126

Piché, J., Hutchings, J. A., & Blanchard, W. (2008). Genetic variation in threshold reaction norms for alternative reproductive tactics in male Atlantic salmon, Salmo salar. Proceedings of the Royal Society B: Biological Sciences, 275(1642), 1571–1575. 10.1098/rspb.2008.0251

Prince, D. J., O’Rourke, S. M., Thompson, T. Q., Ali, O. A., Lyman, H. S., Saglam, I. K., Hotaling, T. J., Spidle, A. P., & Miller, M. R. (2017). The evolutionary basis of premature migration in Pacific salmon highlights the utility of genomics for informing conservation. SCIENCE ADVANCES, 3.

Prokkola, J. M., Åsheim, E. R., Morozov, S., Bangura, P., Erkinaro, J., Ruokolainen, A., Primmer, C. R., & Aykanat, T. (2022). Genetic coupling of life-history and aerobic performance in Atlantic salmon. Proceedings of the Royal Society B: Biological Sciences, 289(1967), 20212500. 10.1098/rspb.2021.2500

R Core Team. (2024). R: A language and environment for statistical computing. R Foundation for Statistical Computing [Computer software]. R Foundation for Statistical Computing. https://www.R-project.org/

Raunsgard, A., Persson, L., Czorlich, Y., Ugedal, O., Thorstad, E. B., Karlsson, S., Fiske, P., & Bolstad, G. H. (2023). Variation in phenotypic plasticity across age-at-maturity genotypes in wild Atlantic salmon. Molecular Ecology, n/a. 10.1111/mec.17229

Reid, J. E., & Chaput, G. (2012). Spawning history influence on fecundity, egg size, and egg survival of Atlantic salmon (Salmo salar) from the Miramichi River, New Brunswick, Canada. ICES Journal of Marine Science ICES, 69(9), 1678–1685. 10.1038/278097a0

Roff, D. A. (2011). Genomic insights into life history evolution. In T. Flatt & A. Heyland (Eds.), Mechanisms of Life History Evolution (pp. 11–25). Oxford University Press. 10.1093/acprof:oso/9780199568765.003.0002

Roff, D. A., & Emerson, K. (2006). Epistasis and Dominance: Evidence for Differential Effects in Life-History Versus Morphological Traits. Evolution, 60(10), 1981–1990. 10.1111/j.0014-3820.2006.tb01836.x

Rowe, D. K., Thorpe, J. E., & Shanks, A. M. (1991). Role of Fat Stores in the Maturation of Male Atlantic Salmon (*Salmo salar*) Parr. Canadian Journal of Fisheries and Aquatic Sciences, 48(3), 405–413. 10.1139/f91-052

Säisä, M., Koljonen, M.-L., Gross, R., Nilsson, J., Tähtinen, J., Koskiniemi, J., & Vasemägi, A. (2005). Population genetic structure and postglacial colonization of Atlantic salmon (*Salmo salar*) in the Baltic Sea area based on microsatellite DNA variation. Canadian Journal of Fisheries and Aquatic Sciences, 62(8), 1887–1904. 10.1139/f05-094

Shearer, K. D., & Swanson, P. (2000). The effect of whole body lipid on early sexual maturation of 1 + age male chinook salmon (Oncorhynchus tshawytscha). Aquaculture, 190(3–4), 343–367. 10.1016/S0044-8486(00)00406-3

Sinclair-Waters, M., Ødegård, J., Korsvoll, S. A., Moen, T., Lien, S., Primmer, C. R., & Barson, N. J. (2020). Beyond large-effect loci: Large-scale GWAS reveals a mixed large-effect and polygenic architecture for age at maturity of Atlantic salmon. Genetics Selection Evolution, 52(1), 1–11. 10.1186/s12711-020-0529-8

Sinclair-Waters, M., Piavchenko, N., Ruokolainen, A., Aykanat, T., Erkinaro, J., & Primmer, C. R. (2022). Refining the genomic location of single nucleotide polymorphism variation affecting Atlantic salmon maturation timing at a key large-effect locus. Molecular Ecology, 31(2), 562–570. 10.1111/mec.16256

Stan Development Team. (2020). RStan: The R interface to Stan (Version 2.19.3) [Computer software]. http://mc-stan.org/

Stearns, S. C. (2000). Life history evolution: Successes, limitations, and prospects. Naturwissenschaften, 87(11), 476–486. 10.1007/s001140050763

Taranger, G. L., Carrillo, M., Schulz, R. W., Fontaine, P., Zanuy, S., Felip, A., Weltzien, F. A., Dufour, S., Karlsen, Ø., Norberg, B., Andersson, E., & Hansen, T. (2010). Control of puberty in farmed fish. General and Comparative Endocrinology, 165(3), 483–515. 10.1016/j.ygcen.2009.05.004

Thorpe, J. E., Mangel, M., Metcalfe, N. B., & Huntingford, F. A. (1998). Modelling the proximate basis of salmonid life-history variation, with application to Atlantic salmon, Salmo salar L. Evolutionary Ecology, 12(5), 581–599. 10.1023/A:1022351814644

Thorpe, J. E., Miles, M. S., & Keay, D. S. (1984). Developmental rate, fecundity and egg size in Atlantic salmon, Salmo salar L. Aquaculture, 43(1–3), 289–305. 10.1016/0044-8486(84)90030-9

Thorstad, E. B., Whoriskey, F., Rikardsen, A. H., & Aarestrup, K. (2010). Aquatic Nomads: The Life and Migrations of the Atlantic Salmon. In Ø. Aas, S. Einum, A. Klemetsen, & J. Skurdal (Eds.), Atlantic Salmon Ecology (pp. 1–32). Wiley-Blackwell. 10.1002/9781444327755.ch1

Tréhin, C., Rivot, E., Lamireau, L., Meslier, L., Besnard, A. L., Gregory, S. D., & Nevoux, M. (2021). Growth during the first summer at sea modulates sex-specific maturation schedule in Atlantic salmon. Canadian Journal of Fisheries and Aquatic Sciences, 78(6), 659–669. 10.1139/cjfas-2020-0236

Tréhin, C., Rivot, E., Santanbien, V., Patin, R., Gregory, S. D., Lamireau, L., Marchand, F., Beaumont, W. R. C., Scott, L. J., Hillman, R., Besnard, A.-L., Boisson, P.-Y., Meslier, L., King, A. R., Stevens, J. R., & Nevoux, M. (2024). A multi-population approach supports common patterns in marine growth and maturation decision in Atlantic salmon (Salmo salar L.) from southern Europe. Journal of Fish Biology, 104(1), 125–138. 10.1111/jfb.15567

Vehtari, A., Gabry, J., Magnusson, M., Yao, Y., Bürkner, P.-C., Paananen, T., Gelman, A., Goodrich, B., Piironen, J., Nicenboim, B., & Lindgren, L. (2024). loo: Efficient Leave-One-Out Cross-Validation and WAIC for Bayesian Models (Version 2.7.0) [Computer software]. https://mc-stan.org/loo/

Vehtari, A., Gelman, A., & Gabry, J. (2017). Practical Bayesian model evaluation using leave-one-out cross-validation and WAIC. Statistics and Computing, 27(5), 1413–1432. 10.1007/s11222-016-9696-4

Vollset, K. W., Urdal, K., Utne, K., Thorstad, E. B., Sægrov, H., Raunsgard, A., Skagseth, Ø., Lennox, R. J., Østborg, G. M., Ugedal, O., Jensen, A. J., Bolstad, G. H., & Fiske, P. (2022). Ecological regime shift in the Northeast Atlantic Ocean revealed from the unprecedented reduction in marine growth of Atlantic salmon. Science Advances, 8(9), eabk2542. 10.1126/sciadv.abk2542

White, J., Fitzgerald, C., Gargan, P., de Eyto, E., Millane, M., Chaput, G., Boylan, P., Crozier, W. W., Doherty, D., Kennedy, B., Lawler, I., Lyons, D., Marnell, F., McGinnity, P., O’Higgins, K., Roche, W. K., Maxwell, H., & Ó Maoiléidigh, N. (2023). Incorporating conservation limit variability and stock risk assessment in precautionary salmon catch advice at the river scale. ICES Journal of Marine Science, 80(4), 803–822. 10.1093/icesjms/fsad006

Wickham, H., Averick, M., Bryan, J., Chang, W., McGowan, L., François, R., Grolemund, G., Hayes, A., Henry, L., Hester, J., Kuhn, M., Pedersen, T., Miller, E., Bache, S., Müller, K., Ooms, J., Robinson, D., Seidel, D., Spinu, V., … Yutani, H. (2019). Welcome to the Tidyverse. Journal of Open Source Software, 4(43), 1686. 10.21105/joss.01686

Wilson, A. J., Réale, D., Clements, M. N., Morrissey, M. M., Postma, E., Walling, C. A., Kruuk, L. E. B., & Nussey, D. H. (2010). An ecologist’s guide to the animal model. Journal of Animal Ecology, 79(1), 13–26. 10.1111/j.1365-2656.2009.01639.x

Woronik, A., Tunström, K., Perry, M. W., Neethiraj, R., Stefanescu, C., Celorio-Mancera, M. de la P., Brattström, O., Hill, J., Lehmann, P., Käkelä, R., & Wheat, C. W. (2019). A transposable element insertion is associated with an alternative life history strategy. Nature Communications, 10(1), 5757. 10.1038/s41467-019-13596-2

Yao, Y., Vehtari, A., Simpson, D., & Gelman, A. (2018). Using Stacking to Average Bayesian Predictive Distributions (with Discussion). Bayesian Analysis, 13(3). 10.1214/17-BA1091

